# Sleep pressure accumulates in a voltage-gated lipid peroxidation memory

**DOI:** 10.1101/2024.02.25.581768

**Authors:** H. Olof Rorsman, Max A. Müller, Patrick Z. Liu, Laura Garmendia Sanchez, Anissa Kempf, Stefanie Gerbig, Bernhard Spengler, Gero Miesenböck

## Abstract

Voltage-gated potassium (K_V_) channels contain cytoplasmic β-subunits whose aldo-keto reductase activity is required for the homeostatic regulation of sleep. Here we show that Hyperkinetic, the β-subunit of the K_V_1 channel Shaker in *Drosophila*, forms a dynamic lipid peroxidation memory. Information is stored in the oxidation state of Hyperkinetic’s nicotinamide adenine dinucleotide phosphate (NADPH) cofactor, which changes when lipid-derived carbonyls, such as 4-oxo-2-nonenal or an endogenous analog generated by illuminating a membrane-bound photosensitizer, abstract an electron pair. NADP^+^ remains locked in the active site of K_V_β until membrane depolarization permits its release and replacement with NADPH. Sleep-inducing neurons use this voltage-gated oxidoreductase cycle to encode their recent lipid peroxidation history in the collective binary states of their K_V_β-subunits; this biochemical memory influences—and is erased by—spike discharges driving sleep. The presence of a lipid peroxidation sensor at the core of homeostatic sleep control suggests that sleep protects neuronal membranes against oxidative damage. Indeed, brain phospholipids are depleted of vulnerable polyunsaturated fatty acyl chains after enforced waking, and slowing the removal of their carbonylic breakdown products increases the demand for sleep.

## Introduction

The pore-forming α-subunits of voltage-gated potassium channels of the K_V_1 and K_V_4 families partner with cytoplasmic β-subunits^1-5^ whose sequences exhibit puzzling similarity with aldo-keto reductases^6,7^—enzymes that reduce carbonyls to alcohols via the coupled oxidation of an NADPH cofactor. The isolated β-subunits show weak reductase activity toward a range of model aldehydes *in vitro*^8,9^, relying on NADPH as the electron donor, but whether, on which native carbonyls, and to what end the assembled K_V_ channel catalyzes similar reactions *in vivo* is unknown. K_V_β’s exceptionally firm grip on its cofactor^10^, which chokes catalysis, deepens the mystery: why would an ion channel be shackled to what appears to be a subpar enzyme?

A hint at a possible answer has come from studies in *Drosophila*, where both the K_V_1 family member Shaker^11,12^ and its β-subunit Hyperkinetic^7^ are needed to sustain normal levels of sleep^13,14^. The sleep-regulatory function of the channel complex has been mapped to a handful of sleep-control neurons whose axonal projections target the dorsal fan-shaped body in the central brain^15,16^ (dFBNs). Sleep need is encoded in the electrical activity of these neurons^17^, which fluctuates—in part^16^—because Hyperkinetic modulates the inactivation kinetics of the Shaker current^18^. During waking, electrons leaking from the saturated transport chains of the inner mitochondrial membrane produce superoxide and other reactive oxygen species (ROS), which convert the K_V_β pool to the NADP^+^-bound form^18^. This prolongs the inactivation time constant of the associated potassium conductance^8,18-20^, strengthens the repolarizing force that restores the resting membrane potential after each spike, and so enables dFBNs to fire at higher rates^18,21^—precisely the changes required to induce sleep^17,22^.

Although the source (the mitochondrial electron transport chain) and the receiver (Hyperkinetic in complex with Shaker) of the sleep-promoting redox signal are known^18^, the mode of communication between mitochondria and potassium channels remains undefined. K_V_β-bound NADPH is an unlikely direct target of ROS, not only because radical-induced hydrogen abstraction (which involves a single electron transfer^23,24^) will not produce NADP^+^ (which would require the loss of two electrons). As ROS spread from the inner mitochondrial membrane^25^, they will encounter many potential reaction partners before reaching Hyperkinetic at the cell surface. Among the most abundant and vulnerable ROS targets in the immediate vicinity of their site of origin are the polyunsaturated fatty acyl chains (PUFAs) of membrane lipids, whose peroxidation and subsequent fragmentation into carbonyls can create chemical functionality fit for the active site of an aldo-keto reductase^23,24,26,27^. In the crystal structure of the mammalian K_V_1.2–β2 channel complex, the substrate binding pocket is lined with hydrophobic residues and filled with unresolved electron density^4^, as would be expected if a diverse group of lipid precursors disintegrated into a heterogeneous mix of apolar ligands. Recombinant K_V_β1 and K_V_β2 reduce synthetic analogs of lipid peroxidation products, such as 4-oxo-2-nonenal (4-ONE), 1-palmitoyl-2-oxovaleroyl-phosphatidylcholine, or methylglyoxal, in the test tube^8,9^, but turnover is so slow that the impact on the concentrations of these molecules *in vivo* must be minimal. While K_V_β can therefore have no plausible role in the enzymatic clearance of toxic carbonyls, the very features that seem detrimental or baroque in a catalyst—the protein’s stranglehold on NADP(H) and its linkage to a voltage-gated ion channel—could be essential if the assembly instead functioned as a biochemical memory cell. Imagine that tight binding of NADP(H) causes the redox reaction to pause at the cofactor exchange step. Each β-subunit then records a single exposure to an oxidizing substrate by flipping from the NADPH-to the NADP^+^-bound form and stores this bit of information until NADP^+^ is released and replaced by NADPH. The operational logic resembles that of a single-transistor dynamic random-access memory (DRAM) cell^28^: K_V_β corresponds to the storage capacitor of a DRAM cell; the oxidation state of NADP(H) plays the part of the electric charge on the capacitor; and the (low) basal reaction rate is equivalent to the leakage of charge from the capacitor, which gives the memory a finite lifetime that requires periodic refreshment^28^. The analogy would be complete if, akin to the voltage across the transistor that gates access to the storage capacitor in a DRAM chip^28^, the membrane potential across the voltage sensors of the α-subunit switched the NADP(H) binding site of the β-subunit between open and closed conformations.

The present study tests several tenets of this model. We examine the lipids of rested and sleep-deprived brains for signs of oxidative damage; measure the impact on sleep of perturbing the clearance of peroxidized lipids; determine if lipid peroxidation products influence the Shaker current of sleep-control neurons via the active site of Hyperkinetic; and analyze the interplay of voltage sensors and NADP(H) binding sites in the redox regulation of the channel. The results define an autoregulatory loop in which the K_V_ channel population encodes the recent lipid peroxidation history of a neuron in the collective binary states of their β-subunits. This biochemical memory (which we equate to the accumulated sleep pressure) is read and erased during subsequent electrical activity, with the action potential frequency set^18^ by the fraction of K_V_β subunits previously loaded with NADP^+^.

## Results

### A lipidomic fingerprint of sleep loss

Because levels of oxidative stress may differ among tissues, brain regions, or neuron types^18,29^, we collected spatial maps of hundreds of lipids by means of high-resolution scanning microprobe matrix-assisted laser desorption/ionization mass spectrometry imaging (SMALDI-MSI). The lipid maps were overlayed on fluorescence images of 10-µm-thick cryosections in which dFBNs were marked with *R23E10–GAL4*-driven^17^ mCD8::GFP (Fig. 1a). Sections were cut from rested or sleep-deprived brains, and technical triplicates of three biological replicates in each condition were scanned at a lateral resolution of 5 µm × 5 µm.

**Fig. 1.**
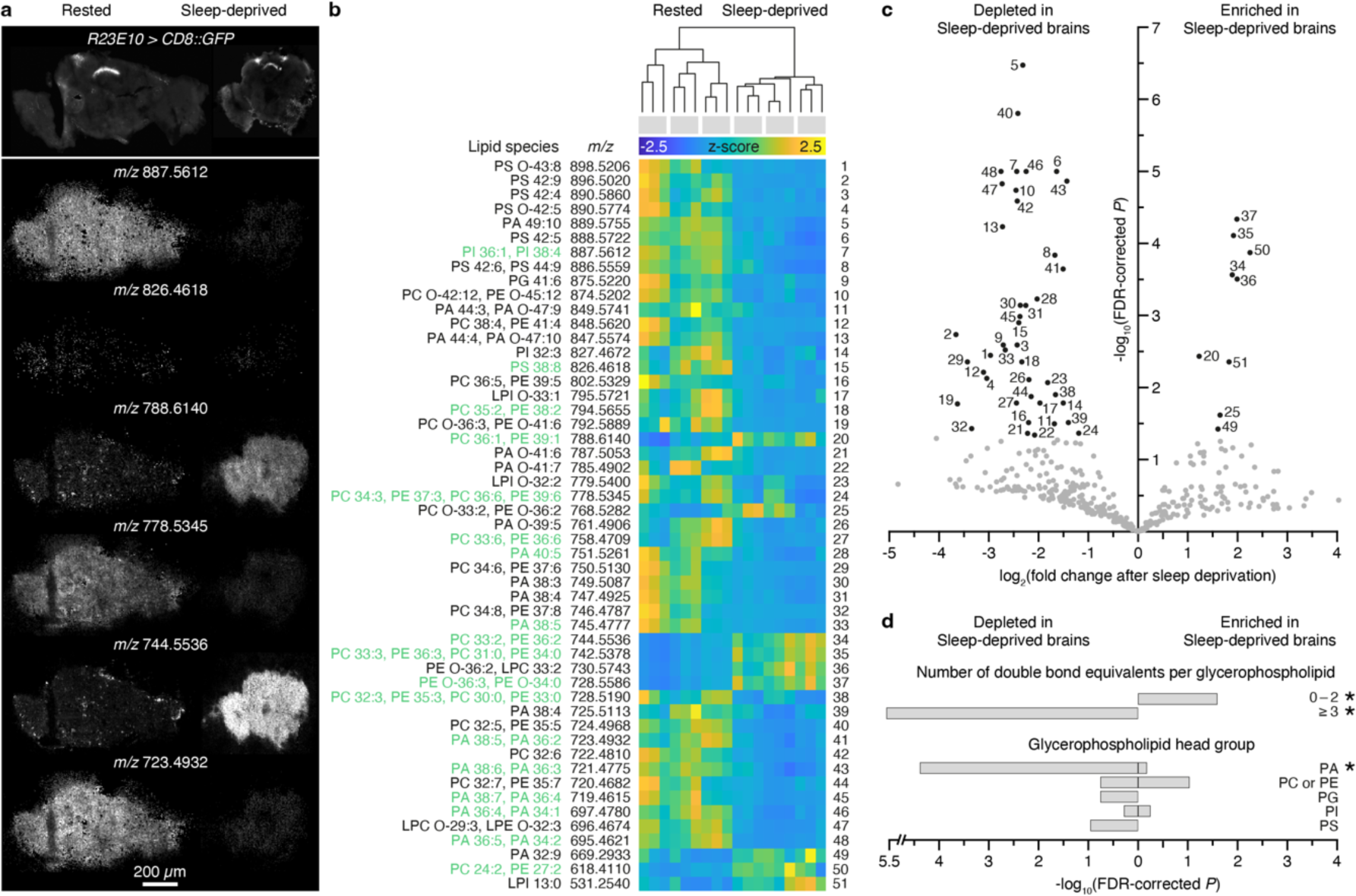
Sleep deprivation depletes brain phospholipids of polyunsaturated fatty acids. **a**, Example fluorescence (top) and positive-ion SMALDI-MS images (bottom) of cryosections containing dFBNs marked with mCD8::GFP. The sections were cut from rested (left) and sleep-deprived brains (right). SMALDI-MS images show, from top to bottom, the spatial distributions of PI 18:2/20:2 (*m/z* 887.5612, [M+Na]^+^), PS 18:3/20:5 (*m/z* 826.4618, [M+Na]^+^), PC 18:0/18:1 (*m/z* 788.6140, [M+H]^+^), PC 18:3/18:3 (*m/z* 778.5345, [M+H]^+^), PE 18:1/18:1 (*m/z* 744.5536, [M+H]^+^), and PA 18:2/20:3 (*m/z* 723.4932, [M+H]^+^). **b**, Hierarchical clustering of rested and sleep-deprived brains according to their glycerophospholipid profiles. Heat maps show the z-scored intensities of *m/z* signals differing with sleep history at an FDR-adjusted *P*<0.05. Lipids detected in MS^2^ fragmentation experiments are annotated in green type in the list of molecular assignments on the left. Each column represents a technical replicate; biological replicates are grouped by gray bars on top. **c**, Volcano plot of sleep history-dependent changes in 380 *m/z* signals annotated as glycerophospholipids. Signals with >2-fold intensity changes and FDR-corrected *P*<0.05 are indicated in black. Numerical labels cross-reference data points to lipid annotations in **b**. **d**, Features overrepresented in the subset of 51 differentially abundant lipids against the background set of all 380 glycerophospholipids. Asterisks indicate significant enrichment scores (FDR-corrected *P*<0.05). Because PC and PE lipids cannot be distinguished by exact mass alone, they are grouped as a single feature. Abbreviations: PA, phosphatidic acid, (L)PC, (lyso)phosphatidylcholine; (L)PE, (lyso)phosphatidylethanolamine; PG, phosphatidylglycerol; (L)PI, (lyso)phosphatidylinositol; PS, phosphatidylserine; O-, alkyl ether linkage.

Samples within each group had tightly correlated lipid profiles, but differences between groups—that is, between the rested and sleep-deprived states—were so stark that sleep histories could be accurately inferred from lipid composition alone; a single principal component captured 85% of the overall variance. Fifty-one among 380 SMALDI-MSI signals annotated as glycerophospholipids and detected exclusively on tissue increased or decreased >2-fold after sleep loss, with a false discovery rate (FDR)-adjusted significance threshold of *P* < 0.05 and little, if any, spatial heterogeneity across the brain (Fig. 1a–c). The identities of 18 of these 51 differentially abundant phospholipids (35%) were confirmed by targeted MS^2^ fragmentation after HPLC separation of a methyl *tert*-butyl ether extract of brain homogenates (Fig. 1a–c). In many cases these analyses also revealed the detailed fatty acid compositions of the parent species (Fig. 1b, c).

Most lipids with high discriminatory power belonged to one of three classes, which form discernible blocks in the clustergram of Fig. 1b. The glycerophospholipids of rested brains carried inositol, serine, ethanolamine, or choline head groups and were enriched in acyl chains with a combined median length of 37.5 carbons and a large degree of unsaturation; the number of double bonds averaged 5.0 ± 2.61 (mean ± s.d.) per lipid, with a median of 5 and a maximum of 12 (Fig. 1b–d). Phospholipids present at higher levels in sleep-deprived brains, by contrast, contained mostly choline and ethanolamine head groups, shorter acyl chains with a combined median length of 33.5 carbons, and many fewer double bonds than those in rested flies; their number averaged 2.0 ± 2.03 (mean ± s.d.) per lipid, with a median of 2 (Fig. 1b– d). The third distinctive lipid class consisted of several species of phosphatidic acid, whose levels declined after sleep deprivation (Fig. 1b–d). Phosphatidic acid occupies a central position in the biosynthetic pathways of all glycerophospholipids^30,31^ and promotes mitochondrial fusion when generated locally by a dedicated phospholipase D (mitoPLD)^32^. A companion study shows that reduced mitoPLD activity in dFBNs causes sleep loss^33^.

The lipidomic fingerprint of sleep-deprived brains indicates that their membranes are depleted of PUFAs, presumably as a consequence of oxidative damage, leaving behind a greater proportion of largely saturated phospholipids (Fig. 1d). The picture during rest is consistent with membrane repair via glycerophospholipid biosynthesis from phosphatidic acid precursors^30,31^ and a reversal of the mitochondrial fragmentation that commonly accompanies periods of oxidative stress^25,34^, including sleep deprivation^33^.

### PUFA-derived carbonyls promote sleep

The peroxidation of membrane lipids begins^23,24^ with the abstraction of a *bis*-allylic hydrogen from a PUFA chain by a radical oxidant such as HOO• (the conjugate acid of O_2-_) or •OH. The resulting lipid radical reacts with O_2_ to form a lipid peroxyl radical, which propagates the chain by abstracting a hydrogen from another PUFA, generating a new lipid radical and a lipid hydroperoxide^23,24,26,27^. The reaction continues until two radicals combine in a termination step. The lipid hydroperoxides produced along the way undergo a series of rearrangements and scissions that give rise to a variety of short-and medium-chain carbonyl breakdown products^23,24,26,27^, including the potential K_V_β substrate^8,9^ 4-ONE.

Operating behind a primary bastion of enzymatic and non-enzymatic antioxidants^35^, soluble short-chain dehydrogenases/reductases, such as carbonyl reductase 1 in mammals^36,37^ and its functional homolog sniffer in *Drosophila*^38,39^, form an outer defensive ring against lipid peroxidation-derived carbonyls. We examined whether breaching and mending these secondary defences would recapitulate the well-documented effects on sleep of pro-and antioxidant manipulations^18,29,40^. Indeed, hemizygous male carriers of the X-linked hypomorphic *sniffer* allele *sni*^1^ showed elevated sleep durations during the day and night (Fig. 2a–c, 1a), due to vastly extended, hyperconsolidated sleep episodes (Supplementary Fig. 1b, c), at an age before widespread neurodegeneration^38^ produced locomotor deficits that could have been mistaken for sleep (Supplementary Fig. 1d). Sleep returned to or below wild-type levels when *sni*^1^ mutants expressed a *UAS–sni* rescue transgene^38^ (Fig. 2c, Supplementary Fig. 1a), and similarly when the alternative oxidase AOX, which shunts electrons from a reduced ubiquinone pool to H_2_O, capped mitochondrial ROS production^18,41^ (Fig. 2a, c), or when the putative carbonyl sensor Hyperkinetic was removed by RNA-mediated interference (RNAi), either pan-neuronally or in dFBNs of *sni*^1^ mutant flies (Fig. 2b, c). These data place lipid peroxidation products downstream of mitochondrial respiration in the signalling chain that terminates on the Hyperkinetic pool of dFBNs to raise the pressure to sleep^18^.

**Fig. 2.**
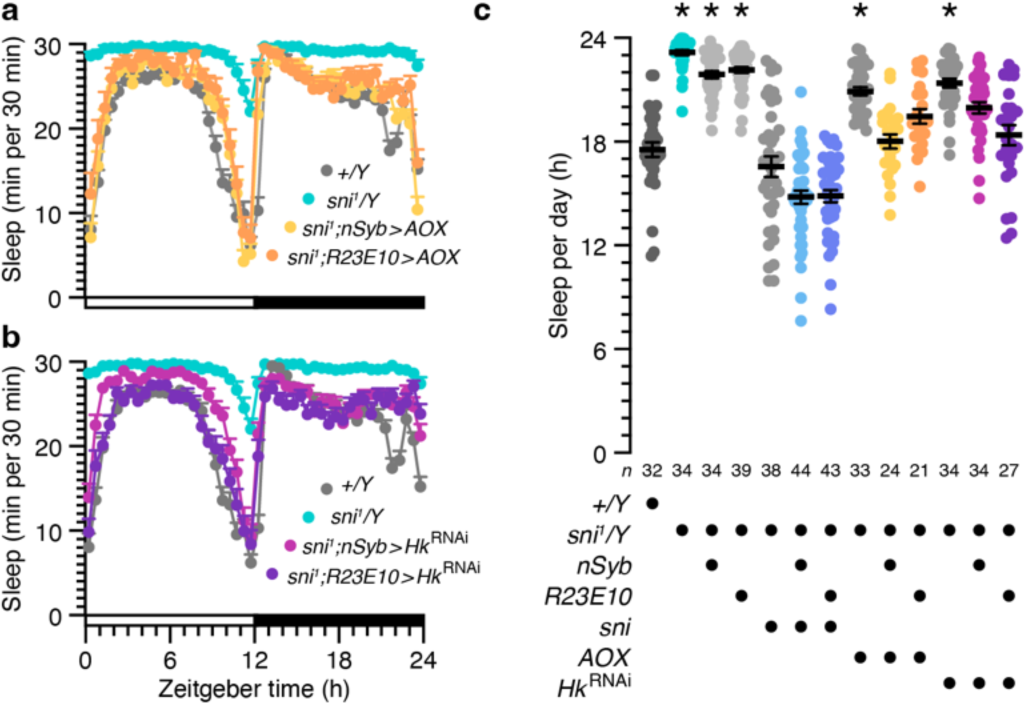
PUFA-derived carbonyls are intermediates in the signalling chain that couples mitochondrial electron transport to sleep. **a**, The *nSyb-GAL4-* or *R23E10-GAL4*-driven expression of AOX in hemizygous *sni^1^* mutant males fully or partially restores wild-type sleep (two-way repeated-measures ANOVA with Holm-Šídák test; sample sizes as in **c**): the sleep profiles of *sni^1^* mutants with pan-neuronal expression of AOX differ from those of *sni^1^* mutants (*P*<0.0001) but not of wild-type flies (*P*=0.0589), while the sleep profiles of *sni^1^*mutants with dFBN-restricted expression of AOX differ from those of both *sni^1^*mutants (*P*<0.0001) and wild-type flies (*P*=0.0007). **b**, *nSyb-GAL4-* or *R23E10-GAL4*-restricted interference with the expression of Hyperkinetic in hemizygous *sni^1^*mutant males partially or fully restores wild-type sleep (two-way repeated-measures ANOVA with Holm-Šídák test; sample sizes as in **c**): the sleep profiles of *sni^1^* mutants with pan-neuronal expression of *Hk*^RNAi^ differ from those of both *sni^1^* mutants (*P*<0.0001) and wild-type flies (*P*<0.0001), while the sleep profiles of *sni^1^* mutants with dFBN-restricted expression of *Hk*^RNAi^ differ from those of *sni^1^* mutants (*P*<0.0001) but not of wild-type flies (*P*=0.1344). **c**, Total sleep in hemizygous males carrying the *sni^1^* allele differs from wild-type (*P*<0.0001; Kruskal-Wallis ANOVA with Dunn’s test) but returns to or below control level if carriers also express sniffer, AOX, or *Hk*^RNAi^ pan-neuronally under the control of *nSyb-GAL4* (sni: *P*=0.1128; AOX: *P*>0.9999; *Hk*^RNAi^: *P*=0.0601) or in dFBNs under the control of *R23E10-GAL4* (sni: *P*=0.1151; AOX: *P*=0.6694; *Hk*^RNAi^: *P*>0.9999). Note that the expression of the *UAS-sni* transgene appears leaky, as the sleep phenotype of *sni^1^*mutants is rescued in the absence of a *GAL4* driver (*P*>0.9999). Asterisks indicate significant differences (*P*<0.05) from wild-type in planned pairwise comparisons. Data are means ± s.e.m. *n*, number of flies. For statistical details see Supplementary Table 1.

### A redox memory of lipid peroxidation

To determine whether lipid-derived carbonyls could alter the oxidation state of Hyperkinetic’s cofactor, we obtained whole-cell voltage-clamp recordings from dFBNs marked with *R23E10-GAL4*-driven mCD8::GFP and relied on the bi-exponential inactivation kinetics of *I*_A_ as our estimate of the NADP^+^:NADPH ratio of a cell’s Hyperkinetic population^18^ (Supplementary Fig. 2). As in previous experiments^18^, we calibrated our measurements with the help of plasma membrane-anchored miniSOG^42^. The exposed flavin mononucleotide chromophore of this light-oxygen-voltage-sensing (LOV) domain protein transfers its excitation energy during blue illumination efficiently to O_2_, producing singlet oxygen (^1^O_2_) which—presumably indirectly—shifts the channel population to the NADP^+^-bound form and induces sleep^18^. The oxidation of the cofactor was detected as an increase in the A-type current’s fast and slow inactivation time constants after 9 minutes of blue light exposure, from initial mean values of 5.8 and 35 ms to final averages of 8.2 and 59 ms (Fig. 3a). When the membrane potential was clamped at –80 mV, *τ*_fast_ and *τ*_slow_ stayed stably elevated for 20 minutes after the light-driven ^1^O_2_ generation stopped (Fig. 3a, b), consistent with a negligible rate of spontaneous NADP^+^ exchange^8,10,20^ that allows Hyperkinetic to retain a memory of an earlier encounter with an oxidizing substrate, which itself is short-lived (intracellular half-life^26^ of lipid-derived aldehydes <4 s).

**Fig. 3.**
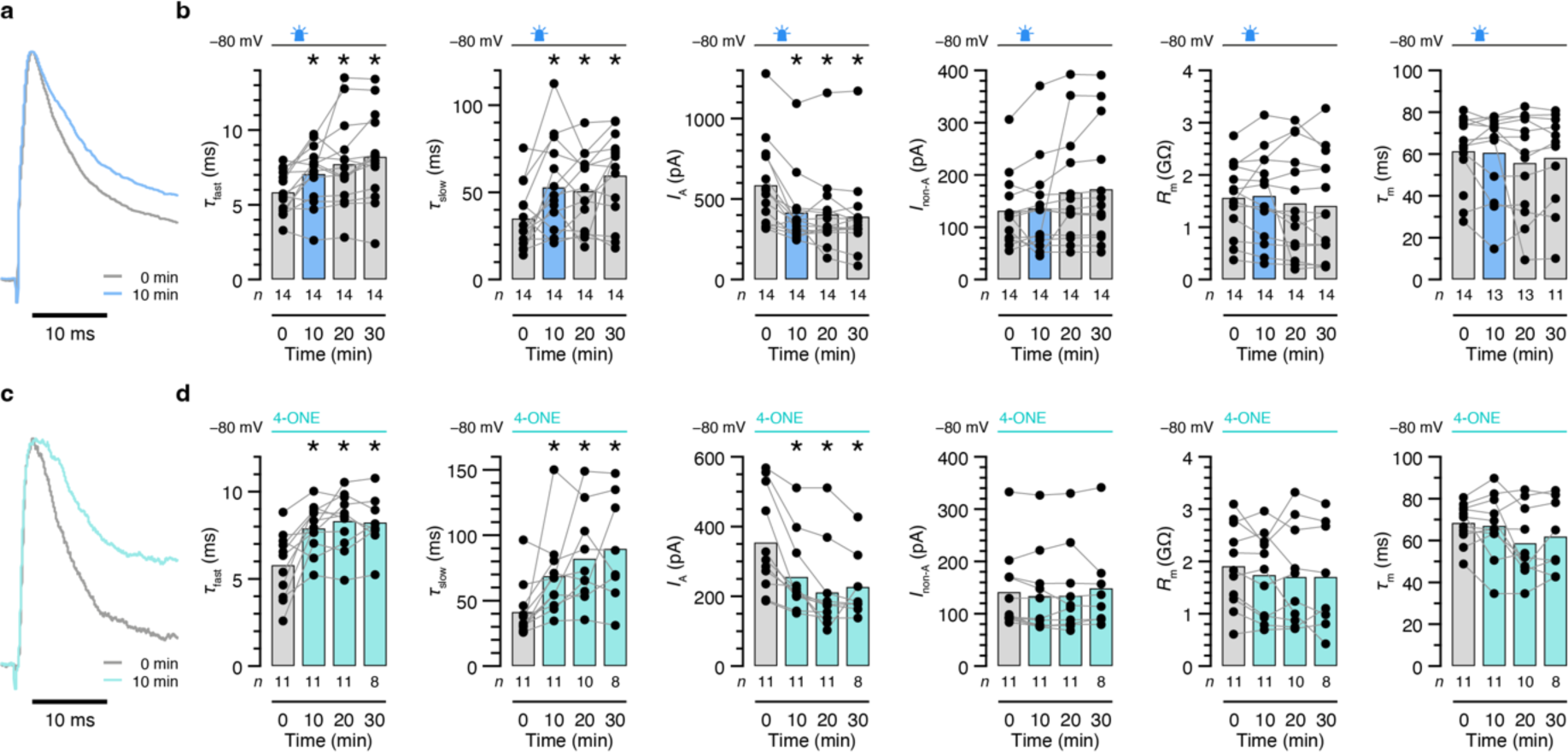
ROS and PUFA-derived carbonyls imprint a memory on the inactivation kinetics of *I*_A_. **a**, **b**, dFBNs expressing miniSOG were held at –80 mV, except during the voltage protocols required to measure *I*_A_. A 9-minute exposure to blue light between the 0-and 10-minute time points (**b**, blue shading) increases the fast and slow inactivation time constants of *I*_A_ stably above their pre-illumination baselines (**b**, *τ*_fast_: *P*=0.0133; *τ*_slow_: *P*=0.0041; repeated-measures ANOVA; examples of peak-normalized *I*_A_ evoked in the same dFBN by voltage steps to +30 mV in **a**). The amplitude of *I*_A_ runs down during the course of the recording (**b**, *P*<0.0001; repeated-measures ANOVA); *I*_non-A_, input resistances (*R*_m_), and membrane time constants (*τ*_m_) remain unchanged (**b**, *P≥*0.0971; repeated-measures ANOVA or mixed-effects model). **c**, **d**, dFBNs were held at –80 mV, except during the voltage protocols required to measure *I*_A_. The inclusion of 50 µM 4-ONE in the intracellular solution (**d**, turquoise shading) increases the fast and slow inactivation time constants of *I*_A_ stably above the baselines measured immediately after break-in (**d**, *τ*_fast_: *P*=0.0015; *τ*_slow_: *P*=0.0010; mixed-effects model; examples of peak-normalized *I*_A_ evoked in the same dFBN by voltage steps to +30 mV in **c**). The amplitude of *I*_A_ runs down during the course of the recording (**d**, *P*=0.0024; mixed-effects model); *I*_non-A_, input resistances (*R*_m_), and membrane time constants (*τ*_m_) remain unchanged (**d**, *P*≥0.0783; mixed-effects model). Columns, population averages; dots, individual cells; *n*, number of cells; asterisks, significant differences (*P*<0.05) relative to baseline in planned comparisons by Holm-Šídák test. For statistical details see Supplementary Table 1.

In a direct test of the idea that PUFA-derived carbonyls are prominent among these substrates, we filled dFBNs through the patch pipette with the synthetic lipid peroxidation products 4-ONE or 4-hydroxynonenal (4-HNE)^24,26,27^. Due to their inherent reactivity and membrane-permeability, the equilibration of these carbonyls within the neuronal arbor was governed by complex reaction–diffusion kinetics that made their concentration profiles difficult to predict^27^ and, in all likelihood, neither spatially uniform nor temporally stationary during the course of a recording. 4-ONE and 4-HNE are estimated (with large uncertainty) to be present in cells in the low to sub-micromolar range under basal conditions but reach millimolar concentrations during periods of oxidative stress^26^. Although 4-HNE is viewed as a useful marker of lipid peroxidation because monoclonal antibodies can detect its protein adducts^43^, mammalian K_V_β2 *in vitro* shows detectable catalytic activity only toward 4-ONE^9^. If the substrate preferences of *Drosophila* Hyperkinetic were similar, 4-HNE could serve as an ideal control to distinguish effects due to the enzymatic conversion of reactive carbonyls from those potentially caused by indiscriminate protein modification^27^.

Comparisons of *I*_A_ inactivation kinetics immediately after break-in and 10 minutes later revealed a clear slowing of the fast and slow time constants, with effect sizes similar to those after the miniSOG-driven photogeneration of ROS (Fig. 3c, d). Changes were seen only in dFBNs loaded with 50 µM 4-ONE; 200 µM 4-HNE, the addition of 0.15% methyl acetate vehicle to the intracellular solution, or the passage of time alone had no effect (Fig. 4, Supplementary Fig. 3). When the cells were held at –80 mV in 4-ONE for extended periods, the inactivation time constants completed much of their climbs to higher plateaux within the first 10 minutes and remained there for the rest of the recordings (Fig. 3d). Because each neuron in this experimental configuration was connected to a practically infinite reservoir of 4-ONE, however, the persistent slowing of inactivation could reflect continuous turnover of substrate rather than a lasting switch in the oxidation state of the cofactor; it can therefore not speak as unequivocally to the longevity of the biochemical memory as the stable elevation of *τ*_fast_ and *τ*_slow_ in miniSOG-expressing dFBNs after a finite light exposure can (Fig. 3b).

**Figure 4.**
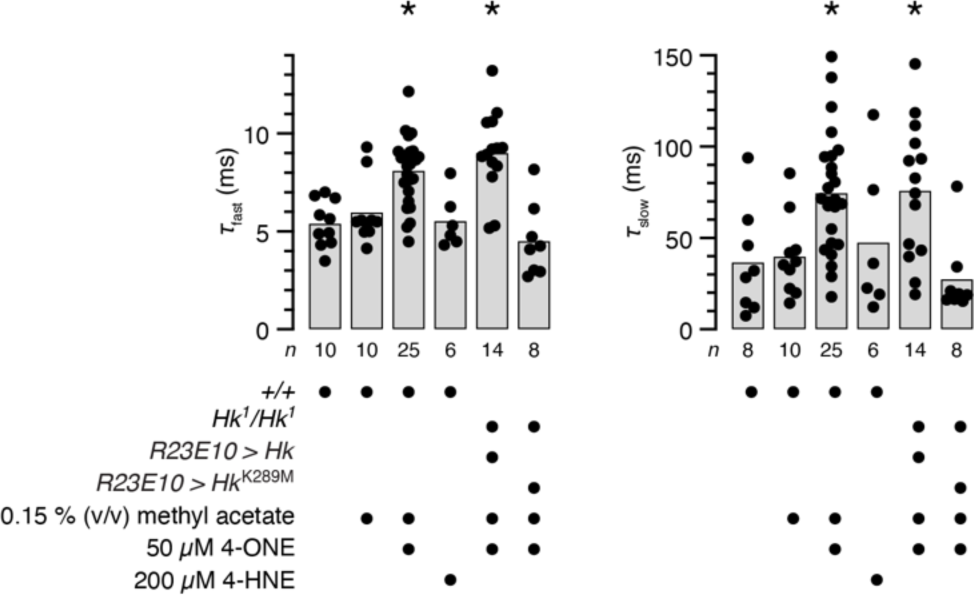
Memory storage requires catalytically active K_V_β and discriminates between 4-ONE and 4-HNE. dFBNs were held at –80 mV, except during the voltage protocols required to measure *I*_A_. At 10 minutes after break-in, the inclusion of 50 µM 4-ONE, but not of 200 µM 4-HNE, in the intracellular solution increases the fast and slow inactivation time constants of *I*_A_ above control levels (*+/+*) in wild-type neurons and neurons expressing a *R23E10-GAL4*-driven *Hk* rescue transgene in a homozygous *Hk^1^* mutant background (*τ*_fast_: *P*<0.0001; *τ*_slow_: *P*=0.0003; Kruskal-Wallis ANOVA). Columns, population averages; dots, individual cells; *n*, number of cells; asterisks, significant differences (*P*<0.05) relative to control levels (*+/+*) in planned comparisons by Dunn’s test. For statistical details see Supplementary Table 1.

Membrane resistances, membrane time constants, and the amplitudes of the non-A-type potassium current remained approximately constant over the course of 30 minutes, but the magnitude of *I*_A_ slowly declined (Fig. 3d). This trend likely reflects closed-state inactivation^44^ rather than a gradual loss of voltage control over a portion of the channels before a rise in access resistance would have prompted us to terminate the recording: series resistances stayed within stable limits for 30 minutes, irrespective of the presence of 4-ONE or changes in command potential or the inactivation kinetics of *I*_A_ (Supplementary Fig. 4a, b), but the steady-state half-inactivation voltages drifted toward more hyperpolarized potentials^44^ (Supplementary Fig. 4c). Because the same slow rundown of *I*_A_ was observed also in the absence of 4-ONE (Supplementary Fig. 3), after miniSOG stimulation (Fig. 3b), and in homozygous *Hyperkinetic* null mutants (see below), the effect cannot be explained by a direct irreversible 4-ONE hit on the β-subunit.

For the most stringent proof that 4-ONE altered the Shaker current via its reduction at the active site of K_V_β (as opposed to an off-target modification on the channel or elsewhere), we expressed transgenes encoding catalytically active or dead Hyperkinetic^45^ under *R23E10-GAL4* control in dFBNs of *Hyperkinetic* null mutant (*Hk*^1^/*Hk*^1^) flies^18^. Infiltrating the Shaker channel with a β-subunit devoid of oxidoreductase activity^8,20,45^ (Hk^K289M^) rendered the fast and slow components of A-type inactivation resistant to 4-ONE, whereas the incorporation of functional K_V_β preserved the channel’s sensitivity (Fig. 4, Supplementary Fig. 5).

While the photogeneration of ROS and the provision of 4-ONE exerted indistinguishable effects in voltage-clamp recordings, current-clamp analyses held a surprise: miniSOG-mediated photooxidation enhanced the spiking response of dFBNs to depolarizing current as expected^18^ (Fig. 5a), whereas 4-ONE did not, even after we corrected for possible differences in input resistance (Fig. 5b). We consider two possible explanations. First, ^1^O_2_ and its reaction products may affect a wider range of molecular targets and cellular processes than 4-ONE and alter spike generation independently of K_V_β. The concerted effect of the active site mutation^8,20,45^ Hk^K289M^ on A-type inactivation and spike frequency, however, makes this an improbable scenario^18^. The second explanation, which we favor, is technical. Because miniSOG is generated biosynthetically, it will easily pervade locations (such as the initial segment and distal branches of dFBN axons) that appear rich in Hyperkinetic but are connected to the soma through long, thin cables (Fig. 5c) and therefore difficult for 4-ONE, with its complex reaction-diffusion kinetics and brief half-life^26,27^, to reach within a short equilibration period. The variable reach of endogenously generated and exogenously supplied oxidants will matter little in measurements of voltage-gated potassium currents, which for space-clamp reasons^46^ are dominated by channels near the somatic recording site (Fig. 5c–e), but will come to the fore in recordings of action potentials if the spike initiation zone lies outside the diffusion distance of 4-ONE.

**Fig. 5.**
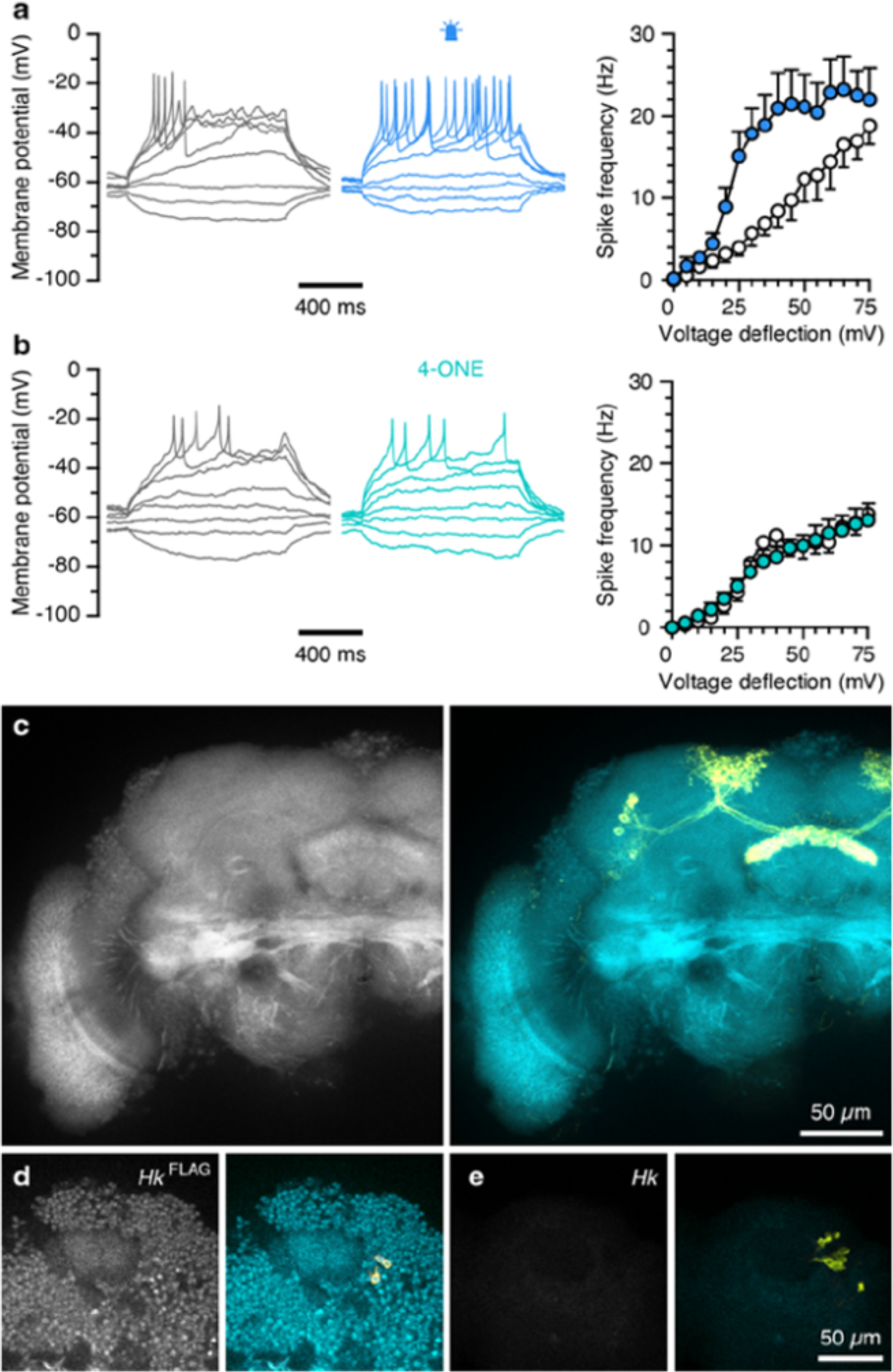
Changes in spike frequency require the conversion of an axonal K_V_β pool to the NADP^+^-bound form. **a**, Example voltage responses to current steps of dFBNs expressing miniSOG after a 9-minute exposure to blue light (blue) or 9 minutes in darkness (gray). In each neuron, the size of the unitary current step was adjusted to produce a 5-mV deflection from a resting potential of –60 ± 5 mV. The blue light exposure steepens the voltage-spike frequency function of illuminated neurons (*n*=6 cells) relative to controls kept in darkness (*n*=7 cells) (illumination effect: *P*=0.0235; current ξ illumination interaction: *P*=0.0008; two-way repeated-measures ANOVA). **b**, Example voltage responses to current steps of dFBNs after 10 minutes of dialysis with 50 µM 4-ONE (turquoise) or methyl acetate vehicle (gray). In each neuron, the size of the unitary current step was adjusted to produce a 5-mV deflection from a resting potential of –60 ± 5 mV. The inclusion of 4-ONE (*n*=12 cells) fails to steepen the voltage-spike frequency function relative to controls (*n*=10 cells) (4-ONE effect: *P*=0.9052; current ξ 4-ONE interaction: *P*=0.7846; two-way repeated-measures ANOVA). **c**–**e**, Summed-intensity projection of a stack of 22 confocal image planes (axial spacing 0.7973 µm) through the fan-shaped body of a fly carrying the *Hk*^FLAG^ allele (**c**) and single confocal image planes through the somatic regions of flies carrying the *Hk*^FLAG^ allele (**d**) or an unmodified *Hk* locus (**e**). Specimens were stained with anti-FLAG antibody (left); native *R23E10-GAL4*-driven mCD8::GFP fluorescence (yellow) is overlaid on the anti-FLAG channel (turquoise) on the right. For statistical details see Supplementary Table 1.

### Voltage changes clear the redox memory

The stability of cofactor binding suggests that each conversion of K_V_β to the NADP^+^-bound state leaves an imprint lasting many minutes (Fig. 3). We equate this imprint—or, more accurately, the imprint on the oxidation state of a dFBN’s Hyperkinetic pool as a whole— with a log of accumulated sleep pressure^18^. As in a digital recording, the binary states of many elementary memory cells thus quantize a continuous variable, with a resolution determined by the number of single-bit units. Because sleep pressure is discharged via the electrical activity of dFBNs^17,22^, action potentials should erase this memory, by releasing NADP^+^ and allowing its replacement with NADPH, whose abundance in the cytoplasm exceeds that of NADP^+^ by at least 40-fold^47^. Such a mechanism would confirm a long-suspected quirk in the enzymatic cycle of K_V_β and offer a rationale for the protein’s association with a voltage-gated ion channel^10,20^.

We tested the prediction that cofactor exchange is voltage-controlled in both of our experimental configurations, using either the photogeneration of ROS by miniSOG (Fig. 6a, b) or the inclusion of 50 µM 4-ONE in the intracellular solution (Fig. 6c, d) to load the K_V_β population with NADP^+^. Following the expected rises of the fast and slow inactivation time constants at 10 minutes after break-in, dFBNs were taken through simulated 20-minute spike trains at 10 Hz under voltage clamp, with each ‘action potential’ consisting of a 3-ms somatic depolarization to +10 mV. Measurements of *τ*_fast_ and *τ*_slow_ after this sequence of voltage steps (that is, at 30 minutes after break-in) showed full reversals of the initial increases driven by miniSOG or 4-ONE (Fig. 6a–d). These reversals were themselves reversible: when dFBNs filled with 4-ONE were held at –80 mV for a further 10 minutes, the large surplus of 4-ONE in the patch pipette once again drove increases in both inactivation time constants (Fig. 6d), whereas a second 9-minute light exposure accomplished the same for miniSOG-expressing cells (Fig. 6b).

**Fig. 6.**
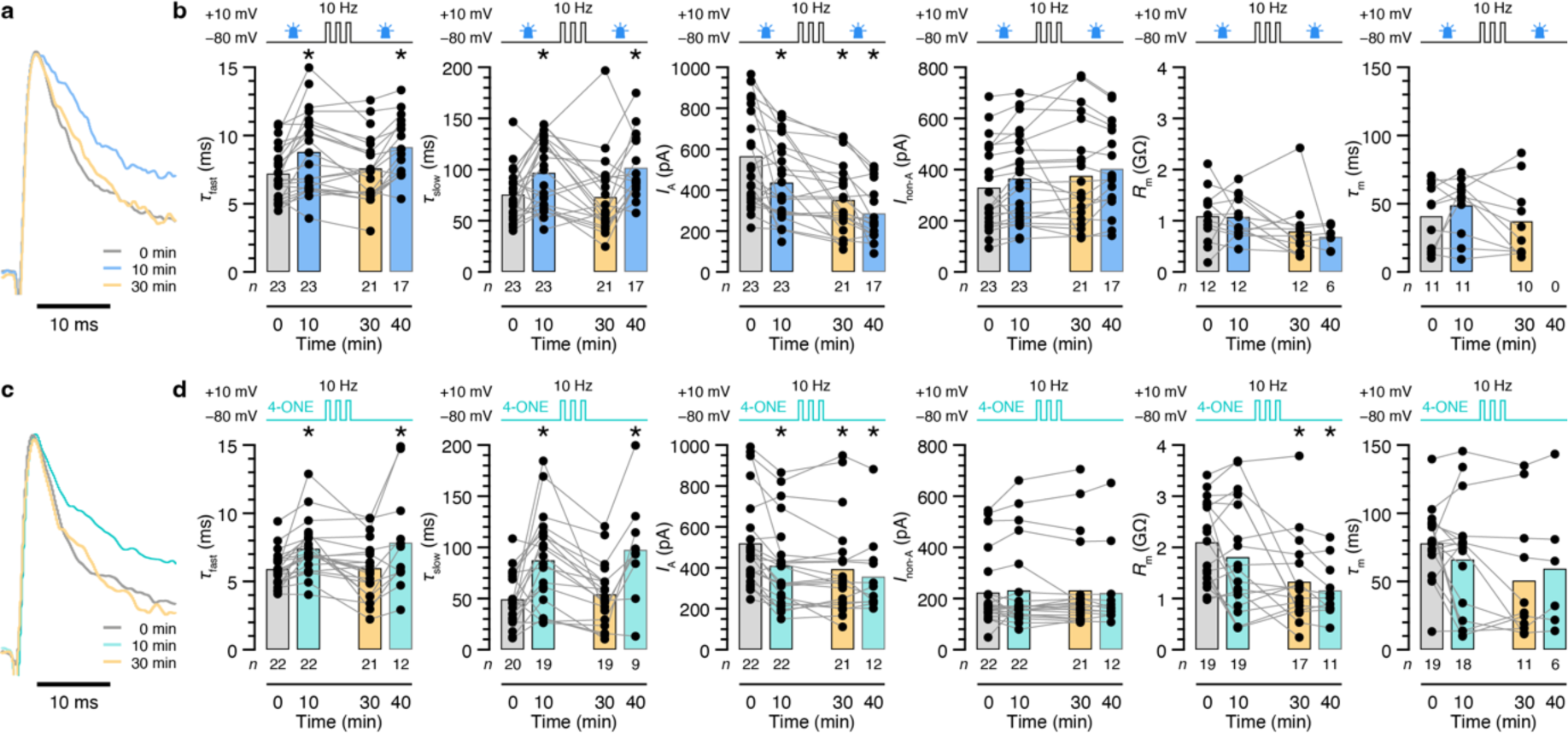
Membrane depolarization erases the lipid peroxidation memory. **a**, **b**, dFBNs expressing miniSOG were held at –80 mV in the intervals of 0–10 and 30–40 minutes (except during the voltage protocols required to measure *I*_A_) and repeatedly step-depolarized to +10 mV (3 ms, 10 Hz) between 10 and 30 minutes. Nine-minute exposures to blue light (between the 0-and 10-minute and the 30-and 40-minute time points) increase the fast and slow inactivation time constants of *I*_A_ above their pre-illumination baselines (**b**, blue vs. gray shading); a series of depolarization steps between 10 and 30 minutes reverses this increase (**b**, yellow shading; *τ*_fast_: *P*<0.0001; *τ*_slow_: *P*=0.0008; mixed-effects model; examples of peak-normalized *I*_A_ evoked in the same dFBN by voltage steps to +30 mV in **a**). The amplitude of *I*_A_ runs down during the course of the recording (**b**, *P*<0.0001; mixed-effects model); *I*_non-A_, input resistances (*R*_m_), and membrane time constants (*τ*_m_) remain unchanged (**b**, *P*≥0.0688; mixed-effects model). **c**, **d**, dFBNs were held at –80 mV in the intervals of 0–10 and 30–40 minutes (except during the voltage protocols required to measure *I*_A_) and repeatedly step-depolarized to +10 mV (3 ms, 10 Hz) between 10 and 30 minutes. The inclusion of 50 µM 4-ONE in the intracellular solution increases the fast and slow inactivation time constants of *I*_A_ above the baseline measured immediately after break-in (**d**, turquoise vs. gray shading); a series of depolarization steps between 10 and 30-minutes counteracts this increase despite the continuous presence of 4-ONE (**d**, yellow shading; *τ*_fast_: *P*=0.0053; *τ*_slow_: *P*=0.0012; mixed-effects model; examples of peak-normalized *I*_A_ evoked in the same dFBN by voltage steps to +30 mV in **c**). The amplitude of *I*_A_ runs down during the course of the recording (**d**, *P*<0.0001; mixed-effects model); input resistances decrease after the series of depolarization steps (*R*_m_ in **d**, *P*=0.0008; mixed-effects model); *I*_non-A_ and membrane time constants (*τ*_m_) remain unchanged (**d**, *P*≥0.1285; mixed-effects model). Columns, population averages; dots, individual cells; *n*, number of cells; asterisks, significant differences (*P*<0.05) relative to baseline in planned comparisons by Holm-Šídák test. For statistical details see Supplementary Table 1.

Occasionally, the reversal protocol pushed the inactivation time constants below their original baselines, suggesting that depolarization dissipated not only the oxidative strain applied by 4-ONE or miniSOG but also the internally sourced pressure already integrated by the channel complex before the experiment began. Consistent with this idea, dFBNs expressing catalytically inactive^8,20,45^ Hk^K289M^, which cannot form a redox memory (Fig. 4, Supplementary Fig. 5a, b), often exhibit the fastest-inactivating A-type currents at baseline^18^ and no modulation by 4-ONE or subsequent voltage changes (Supplementary Fig. 5a, b).

The ability to remember exposures to lipid peroxidation products is an intrinsic property of K_V_1 channels, conserved from insects to mammals, with broad—though not limitless^9^ (Fig. 4)—carbonyl selectivity. When HEK-293 cells coexpressing mouse K_V_1.4 and K_V_β2 were incubated in extracellular medium containing 12 mM methylglyoxal, a membrane-permeable dicarbonyl that serves as an established substrate^9^ for K_V_β2, the fast and slow inactivation time constants of the reconstituted A-type current rose and remained durably elevated for 20 minutes after the removal of methylglyoxal (Supplementary Fig. 6a–c). As in dFBNs expressing Shaker–Hyperkinetic (Fig. 6), the memory was retained if the membrane containing the K_V_1.4–β2 complex was clamped at –80 mV but forgotten during a simulated 20-minute spike train at 10 Hz (Supplementary Fig. 6d, e).

## Discussion

Our experiments suggest that K_V_β subunits are voltage-gated biochemical memories used by neurons and other excitable cells to keep score of lipid peroxidation events. Information is stored in the oxidation state of a nicotinamide molecule bound so tightly that it should perhaps be considered a prosthetic group rather than a cofactor, even though two steps in a stop-and-go redox reaction cycle—hydride transfer and nicotinamide exchange—are used to move data to and from memory. Definitive proof that peroxidized lipids or their breakdown products are endogenous K_V_β substrates would require their co-purification with the native ion channel—a formidable challenge not only because of the expected molecular heterogeneity of these substrates^23,24,26,27^ but also because their binding to K_V_β may be much looser than that of NADP(H); in contrast to the nucleotide binding cleft, which resembles a locked vise, the active site appears wide open in the crystal structure^10^.

Our experiments also suggest, but do not prove beyond doubt, that sleep loss causes widespread lipid peroxidation in the brain. Definitive proof would require a demonstration that peroxidation products accumulate, rather than that polyunsaturated phospholipids are depleted, as we have shown. Most previous attempts to measure lipid peroxidation after sleep loss have focused on a single end product, malondialdehyde^26^, and yielded variable results^48-50^, perhaps because the picture seen through the lens of malondialdehyde is incomplete^43^ or because the thiobarbituric acid assay used for its detection reports tissue oxidizability during analysis rather than pre-existing levels of peroxidized lipids. Our own attempts to quantify endogenous 4-ONE after sleep deprivation succumbed to the double threat of a short-lived, highly reactive analyte^26,27^ with labile, O_2_-sensitive precursors: while SMALDI-MSI could easily detect 100 µM 4-ONE in isolation, the signal vanished when the same quantity of standard was spiked onto a brain section full of endogenous carbonyl-reactive nucleophiles^26,27^ (Supplementary Fig. 7a). Trace amounts of 4-ONE captured by Girard’s reagent during the derivatization of rested, but not sleep-deprived, brains clearly reflected the oxidation of the undepleted PUFA pools of these samples *in vitro* because the *sni*^1^ mutation, which would have raised 4-ONE levels *in vivo*^38,39^, caused no discernible increase at the time of measurement (Supplementary Fig. 7b).

While our interpretation of sleep pressure as mitochondrially-determined^18,33^ lipid peroxidation history demands that sleep-control neurons are equipped to sense and respond to this history, integral redox sensors are a universal feature of K_V_1 (and also some K_V_4) channels^1-5^ in virtually all neurons and many other electrically excitable cells. What could be the purpose of β-subunits in this wider context? Redox control of electrical activity may protect non-renewable cells with high respiratory capacity and extensive membrane systems—such as those of the brain and heart—from oxidative damage if the electron supply to their mitochondria surpasses the demands of ATP synthesis^33^. Depending on where this relief valve opens, the consequences may range from a few extraneous action potentials^21^ (in order to re-balance energy consumption with mitochondrial electron flux) to the induction of sleep^18^. Just as sodium spikes are universal information carriers filled with distinctive meaning by the different neurons that emit them, excitability control by K_V_β may be a general mechanism co-opted by dFBNs for the special purpose of regulating sleep.

## Methods

### Drosophila strains and culture

Flies were reared on media of cornmeal (62.5 g l^-1^), inactive yeast powder (25 g l^-1^), agar (6.75 g l^-1^), molasses (37.5 ml l^-1^), propionic acid (4.2 ml l^-1^), tegosept (1.4 g l^-1^), and ethanol (7 ml l^-1^) on a 12 h light:12 h dark cycle at 25 °C. All electrophysiological and lipidomic analyses were performed on randomly selected females aged 2–6 days post eclosion. Experimental flies were heterozygous for all transgenes and homozygous for either a wild-type or mutant (*Hk^1^*) *Hyperkinetic* allele^51,52^, as stated. The *R23E10-GAL4* driver^17,53^ controlled the expression of the fluorescent label mCD8::GFP in dFBNs, along with an N-myristoylated covalent hexamer (myr-MS6T2) of the singlet oxygen generator miniSOG^54^ or catalytically defective (Hk^K289M^) or functional versions of Hyperkinetic^45^, as indicated.

In behavioral experiments or 4-ONE analyses, hemizygous males carried the X-linked *sni^1^* mutation^38^ and expressed *UAS–sni*^38^, *UAS–AOX*^55^, or *UAS–Hk*^RNAi^ (47805GD)^56^ transgenes, either pan-neuronally^57^ under the control of *nSyb–GAL4* or in dFBNs^17,53^ under the control of *R23E10-GAL4*, as noted.

A *Hyperkinetic* allele encoding an in-frame fusion to an N-terminal FLAG epitope (*Hk*^FLAG^) was created through homology-dependent repair of a CRISPR–Cas9-generated double-strand break (WellGenetics). The FLAG tag was inserted immediately after the initiating methionine of isoforms Hk-PK, Hk-PE, Hk-PL, and Hk-PM and connected to the remainder of the protein via a flexible linker (4 × Gly-Gly-Ser).

### Sleep measurements and sleep deprivation

Females or hemizygous *sni^1^* mutant males^38^ aged 2–5 days were individually inserted into 65-mm glass tubes, loaded into *Drosophila* Activity Monitors (Trikinetics), and housed under 12 h light : 12 h dark conditions. Flies were allowed to adapt to the monitors for a day, and the activity counts during the following two 24-hour periods were averaged. Inactivity periods of >5 minutes were classified as sleep^58,59^ (Sleep and Circadian Analysis MATLAB Program^60^). Immobile flies (<2 beam breaks per 24 h) were manually excluded.

To deprive flies of sleep before lipidomic analyses, a spring-loaded platform stacked with Trikinetics monitors was slowly tilted by an electric motor, released, and allowed to snap back to its original position^61^. The mechanical cycles lasted 10 s and were repeated continuously for 12 h, beginning at zeitgeber time 12.

### SMALDI mass spectrometry imaging

Dissected brains of rested and sleep-deprived flies were placed on PTFE-printed glass slides (Electron Microscopy Sciences), covered with ∼3– 5 µl gelatine (5% w/v in water), and snap frozen for shipping. For sectioning, dissected brains were thawed, suspended in 20 µl 5% gelatine, and transferred to a gelatine plateau created by removing the top half of a frozen block of 5% gelatine in a cryostat (Microm HM 525, ThermoFisher). After allowing the samples to refreeze during 10 minutes in the cryostat chamber, 10 µm thin sections were cut and thaw-mounted onto glass slides. The sections were imaged in fluorescence (BX41, Olympus) and reflected light mode (VHX 5000, Keyence) and stored at –80 °C until further use.

For SMALDI-MSI^62^, the brain sections were thawed in a desiccator and spray-coated with 80 µl of a freshly prepared solution of 2,5-dihydroxybenzoic acid (DHB, Merck) using a SMALDIPrep ultrafine pneumatic spraying system (TransMIT GmbH). The DHB solution contained 60 mg of DHB in 999 µl acetone, 999 µl water, and 2 µl pure trifluoroacetic acid (TFA, Merck). In samples destined for 4-ONE analysis, a chemical derivatization step with Girard’s reagent T (GirT, TCI Chemicals) preceded the application of the DHB matrix^63^. The samples were spray-coated with 35 µl of a freshly prepared solution of 15 mg ml^-1^ GirT in a 7:3 mixture of methanol and water containing 0.2% (v/v) TFA and incubated in a desiccator at room temperature for 2 h. Standards were prepared by applying 5 µl-droplets of a 10-fold dilution series of 4-ONE (Cayman Chemical) in methyl acetate, from 100 µM to 10 nM, onto blank glass slides or slides containing brain sections of rested flies. Standards underwent the same GirT-derivatization and matrix application steps as analytical samples.

A home-built SMALDI-MS imaging ion source based on an AP-SMALDI^5^ AF system (TransMIT GmbH) was coupled to an orbital trapping mass spectrometer (Q Exactive, ThermoFisher). Mass spectra were acquired at a mass resolution of 140,000 in positive-ion mode. A high voltage of 4 kV was applied to the sample holder. The standard pixel size of 5 µm × 5 µm in lipid analyses was increased to 25 µm × 25 µm for 4-ONE measurements to facilitate the detection of low-intensity signals. A single-ion-monitoring (SIM) experiment was performed first for 4-ONE, followed by a full MS scan.

SMALDI-MS images were created in Mirion^64^ (TransMIT GmbH) using a bin width of Δ(*m*/*z*) = 0.004; the images were normalized to total ion charge^65^. A digital mask created from a ubiquitous lipid signal was applied to the measurement area in order to exclude off-tissue pixels, and all images were stitched together in a single file to ensure uniform evaluation. An automatically generated list of all signals found in at least 10 pixels in the stitched file was applied to the separate images to obtain the summed intensity of each signal. Signals were annotated in a bulk search against LIPID MAPS^66^, allowing for [M+H]^+^, [M+Na]^+^, and [M+K]^+^ adducts and selecting the most likely lipid(s) for each measured mass. All annotations with a mass deviation <5 ppm were exported for further validation in HPLC MS^2^ fragmentation experiments.

### HPLC MS^2^ fragmentation

Approximately 1,300 rested and 1,300 sleep-deprived brains were collected in batches of 20–50 per session and snap frozen in plastic tubes. The frozen batches were combined in a glass Potter homogenizer, suspended in 50 µl ice-cold ammonium acetate (0.1% in water, Honeywell), manually homogenized, and transferred into a pre-cleaned Eppendorf tube. Lipids were extracted with 600 µl ice-cold methyl *tert*-butyl ether (MTBE, Sigma-Aldrich) and 150 µl methanol (VWR). After shaking the mixture for 1 h at 4 °C, 200 µl water (VWR) was added, the mixture was shaken for another 10 minutes, and the organic phase was collected after centrifugation for 5 minutes at 1,000 *g*. The aqueous phase was re-extracted using an additional 400 µl MTBE, 120 µl methanol, and 100 µl water. The organic phases from both extraction steps were combined, and the solvent was evaporated under a stream of nitrogen for 30 minutes, leaving ∼700 µg and ∼800 µg of dry extract of rested and sleep-deprived samples, respectively. The extracts were stored at –80°C until further use. An extraction blank was created by performing these steps without brain tissue.

Lipid extracts were thawed, dissolved in 650 µl acetonitrile, 300 µl isopropanol, and 50 µl water (all VWR) in an ultrasonic bath, and separated on a C18 column (100 mm × 2.1 mm, 2.6 µm particle size, 100 Å pore size; Phenomenex Inc.) in an UltiMate 3000 Rapid Separation System (ThermoFisher) coupled to an orbital trapping mass spectrometer (Q Exactive HF-X, ThermoFisher) using a heated electrospray ionization source (HESI II, ThermoFisher). Data-dependent acquisition and MS^2^ fragmentation experiments were based on the inclusion list obtained from SMALDI-MSI annotations, with [M+H] ^+^, [M+Na] ^+^, [M+K] ^+^, and [M+NH_4_]^+^ adducts in positive-ion mode. Since the ionization mechanisms of MALDI and electrospray MS differ, MS^2^ fragmentation of lipid extracts was additionally performed in negative-ion mode, considering [M–H]^-^ and [M+CHO_2_]^-^ adducts, to increase the molecular coverage of SMALDI-MSI hits. Lipids were identified using LipidMatch^67^. All MS^2^-verified lipid annotations were validated by accurate mass and the detection of all fatty acids plus the head group. Only one annotation (PE 27:2) was based on accurate mass and head group detection alone.

### Electrophysiology

Adult female flies aged 2–6 days post eclosion were head-fixed to a custom mount using eicosane (Sigma). Cuticle, trachea, excess adipose tissue, and the perineural sheath were removed to create a small window, and the brain was continuously superfused with extracellular solution equilibrated with 95% O_2_–5% CO_2_ and containing (in mM) 103 NaCl, 3 KCl, 5 TES, 8 trehalose, 10 glucose, 7 sucrose, 26 NaHCO_3_, 1 NaH_2_PO_4_, 1.5 CaCl_2,_ 4 MgCl_2_, pH 7.3, 275 mOsM. GFP-positive cells were visualized on a Zeiss Axioskop 2 FS mot microscope equipped with a 60 ×/1.0 NA water-immersion objective (LUMPLFLN60XW, Olympus) and a pE-300 white LED light source (CoolLED). Borosilicate glass electrodes (9– 11 MΩ) were fabricated on a PC-10 micropipette puller (Narishige) or a DMZ Universal Electrode Puller (Zeitz) and filled with intracellular solution containing (in mM) 10 HEPES, 140 potassium aspartate, 1 KCl, 4 MgATP, 0.5 Na_3_GTP, 1 EGTA, pH 7.3, 265 mOsM. Where indicated, 50 µM 4-ONE or 200 µM 4-hydroxynonenal (4-HNE, Cayman Chemical) were added directly to the intracellular solution. Stock solutions of 4-ONE and 4-HNE were prepared in methyl acetate and ethanol, respectively; vehicle concentrations were not allowed to surpass 0.15% of the total volume after dilution. Recordings were obtained at room temperature with a MultiClamp 700B amplifier, lowpass-filtered at 10 kHz, and sampled at 20 or 50 kHz using Digidata 1440A or 1550B digitizers controlled through pCLAMP 10 or 11 (Molecular Devices). For photostimulation of miniSOG during whole-cell recordings^18^, a 455-nm LED (Thorlabs M455L3) with a mounted collimator lens (Thorlabs ACP2520-A) and T-Cube LED Driver (Thorlabs) delivered 3.5–5 mW cm^-2^ of optical power to the sample. Data were analysed with custom procedures, using the NeuroMatic package (http://neuromatic.thinkrandom.com) in Igor Pro (WaveMetrics).

Whole-cell capacitance compensation and bridge balance were used in voltage-and current-clamp recordings from dFBNs, respectively. Series resistances were monitored but not compensated and allowed to rise at most 20% above baseline—but never beyond 50 MΩ—during a recording. Uncompensated mean series resistances of ∼40 MΩ (Supplementary Fig. 4a) caused predicted voltage errors of ∼16 mV at typical *I*_A_ amplitudes of ∼400 pA (Fig. 3, Fig. 6, Supplementary Fig. 5). Input resistances were calculated from linear fits of the steady-state voltage changes elicited by 1-s steps of hyperpolarizing currents (5-pA increments) from a pre-pulse potential of –60 ± 5 mV. Membrane time constants were estimated by fitting a single exponential to the voltage deflection caused by a hyperpolarizing 5-pA current step lasting 200 ms. Current-frequency curves were determined from voltage responses to a series of depolarizing current steps from a membrane potential of –60 ± 5 mV. To account for variations of input resistances within the dFBN population, the current required to produce a 5-mV hyperpolarizing voltage deflection from a pre-pulse potential of –60 ± 5 mV was used as a cell-specific unitary current step instead of a static 5-pA increment. Spikes were detected by finding minima in the time derivative of the membrane potential trace.

Voltage-clamp experiments on dFBNs were performed in the presence of 1 µM tetrodotoxin (TTX, Tocris) and 200 µM cadmium to block sodium and calcium currents, respectively. Potassium currents were measured by stepping neurons from holding potentials of –10 or – 110 mV for 400 ms to a series of test potentials spanning the range from –100 mV to +30 mV in 10-mV increments^16,18^. Depolarizations from –110 mV produced the sum total of the cell’s potassium currents (*I*_total_, Supplementary Fig. 2a), whereas currents evoked by voltage steps from a holding potential of –10 mV lacked the *I*_A_ (A-type or fast outward) component because voltage-gated potassium channels such as Shaker inactivated (Supplementary Fig. 2b). *I*_A_ was calculated by subtracting this non-A-type component from *I*_total_ (Supplementary Fig. 2c). To determine the fast and slow inactivation time constants^18^, double-exponential functions were fit to the decaying phase of A-type currents elicited by 400-ms steps to +30 mV (Supplementary Fig. 2d). In cases where the fits of slow inactivation time constants were poorly constrained, only the fast inactivation time constants were included in the analysis. Spiking was simulated by 3-ms depolarizing pulses to +10 mV, repeated at 10 Hz for 20 minutes.

Steady-state activation parameters were determined by applying depolarizing 400-ms voltage pulses from holding potentials of –10 or –110 mV; the pulses covered the range from –60 to +60 mV in steps of 10 mV. Linear leak currents were estimated from the slope of the current-voltage relationship at hyperpolarized potentials and subtracted. Steady-state inactivation parameters were obtained with the help of a two-pulse protocol, in which a 300-ms pre-pulse (–120 to +60 mV in 10-mV increments) was followed by a 400-ms test pulse to +30 mV; non-inactivating outward currents, measured from a pre-pulse potential of +10 mV, were subtracted. Peak A-type currents (*I*_A_) were normalized to the maximum current amplitude (*I*_max_) of the respective cell and plotted against the test or pre-pulse potentials (*V*). An estimated liquid junction potential^68^ of 16.1 mV was subtracted post hoc. Curves were fit to the Boltzmann function 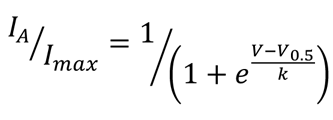 to determine the half-maximal activation and inactivation voltages (*V*_0.5_) and slope factors (*k*).

HEK-293 cells grown in DMEM with 10% fetal bovine serum were transfected (Lipofectamine 3000, ThermoFisher) with a 1:1 mixture of CMV promoter-driven expression vectors encoding mouse K_V_1.4 and a bicistronic mouse K_V_β2–IRES2–EGFP cassette. A carbonyl-reactive residue^69^ (Cys-13) in the N-terminal inactivation peptide of K_V_1.4 was mutated to serine. The growth medium was replaced during whole-cell recordings with extracellular solution containing (in mM) 10 HEPES, 140 NaCl, 5 KCl, 10 glucose, 2 CaCl_2,_ 1 MgCl_2_, pH 7.4. Where indicated, HEK-293 cells were pre-incubated in extracellular solution supplemented with 12 mM methylglyoxal^9^ for 1 h, followed by three washes with methylglyoxal-free solution, before data acquisition. GFP-positive cells were visually targeted with borosilicate glass electrodes (2–3 MΩ) filled with intracellular solution containing (in mM) 10 HEPES, 80 potassium aspartate, 60 KCl, 10 glucose, 2 MgATP, 1 MgCl_2_, 5 EGTA, pH 7.3. Signals were acquired at room temperature with a MultiClamp 700B amplifier, lowpass-filtered at 10 kHz, and sampled at 20 kHz using a Digidata 1440A digitizer controlled through pCLAMP 10 (Molecular Devices). Because untransfected HEK-293 cells lacked voltage-gated conductances (Supplementary Fig. 6a), no channel blockers were present. To determine the fast and slow inactivation time constants, double-exponential functions were fit to the decaying phase of A-type currents elicited by 1-s steps to +30 mV. Spiking was simulated by 3-ms depolarizing pulses to +10 mV, repeated at 10 Hz for 20 minutes. Data were analysed with custom procedures, using the NeuroMatic package (http://neuromatic.thinkrandom.com) in Igor Pro (WaveMetrics).

### Confocal imaging

Dissected brains were fixed for 20 minutes in PBS with 4% (w/v) paraformaldehyde, washed three times for 20 minutes with 0.5% (v/v) Triton X-100 in PBS (PBST), and incubated sequentially at 4 °C in blocking solution (10% goat serum in PBST) overnight, with mouse monoclonal anti-FLAG M2 antibodies (1:500, Sigma) in blocking solution for two days, and with goat anti-Mouse Alexa Fluor 633 antibodies (1:500, ThermoFisher) for one day. The samples were washed five times with blocking solution before and after the addition of the secondary antibody, mounted in Vectashield, and imaged on a Leica TCS SP5 confocal microscope with an HCX IRAPO L 25×/0.95 water immersion objective.

### Quantification and statistical analysis

SMALDI-MSI signal intensities were analysed in LipidSig^70^ and MATLAB (The MathWorks). Global differences between normalized glycerophospholipid intensities in rested and sleep-deprived samples were evaluated by multiple *t* tests with a false discovery rate (FDR)-adjusted *P*<0.05, using the method of Benjamini-Hochberg. Statistical associations with sleep history of user-defined lipid features, such as the indicated double-bond-equivalent ranges or phospholipid head groups, were computed by Fisher’s exact test in LipidSig^70^. Principal component and hierarchical cluster analyses were performed in MATLAB. The list of significantly different signals was exported and re-imported into Mirion to generate SMALDI-MS images for display.

Behavioral and electrophysiological data were analysed in Prism 10 (GraphPad). All null hypothesis tests were two-sided. To control type I errors, *P*-values were adjusted to achieve a joint α of 0.05 at each level in a hypothesis hierarchy; multiplicity adjusted *P*-values are reported in cases of multiple comparisons at one level. Group means or their time courses were compared by paired *t* test, one-or two-way repeated-measures ANOVA, or mixed-effects models in cases where a variable was not measured in all cells at all time points, as indicated in figure legends. Repeated-measures ANOVA and mixed-effect models used the Geisser-Greenhouse correction in all instances except the comparisons of >2 genotypes in Fig. 2a, b and Supplementary Fig. 1a and were followed by planned pairwise analyses with Holm-Šídák’s multiple comparisons test. Where the assumption of normality was violated (as indicated by D’Agostino-Pearson test), group means were compared by Mann-Whitney test, Wilcoxon test, or Kruskal-Wallis ANOVA followed by Dunn’s multiple comparisons test.

## Acknowledgments

We thank L. Ballenberger and C. Hartmann for help with dissections and B. Ganetzky, T. Holmes, H. Jacobs, J. Ng, G. Rubin, S. Schneuwly, J. Simpson, the Bloomington Stock Center, and the Vienna *Drosophila* Resource Center for flies. This work was supported by grants from the European Research Council (832467) and the UK Medical Research Council (MR/V013238/1) to G.M., and from the German Research Council (Sp314/23-1, INST 162/500-1 FUGG) and the Hessian Ministry of Science and Education (LOEWE Center DRUID) to B.S.; H.O.R. and L.G.S. received doctoral training fellowships from Wellcome and La Caixa, respectively; M.A.M. was supported by a Kekulé fellowship from the German Fonds der Chemischen Industrie; P.Z.L. was a Marshall Scholar; and A.K. held postdoctoral fellowships from the Swiss National Science Foundation and EMBO.

## Author contributions

H.O.R. performed all electrophysiological and behavioral experiments on flies and M.A.M. all lipidomic analyses, under the supervision of S.G. and B.S., on material prepared by L.G.S., H.O.R., and A.K.; P.Z.L. characterized *Hk*^FLAG^ flies and K_V_ currents in HEK-293 cells. B.S. designed the SMALDI-MSI methodology and instrumentation. G.M. devised and directed the research and wrote the paper.

## Competing interests

M.A.M. and S.G. are employees and B.S. is a consultant of TransMIT GmbH. All other authors declare no competing interests.

Correspondence and requests for materials should be addressed to G.M.

---

**Supplementary Fig. 1.**
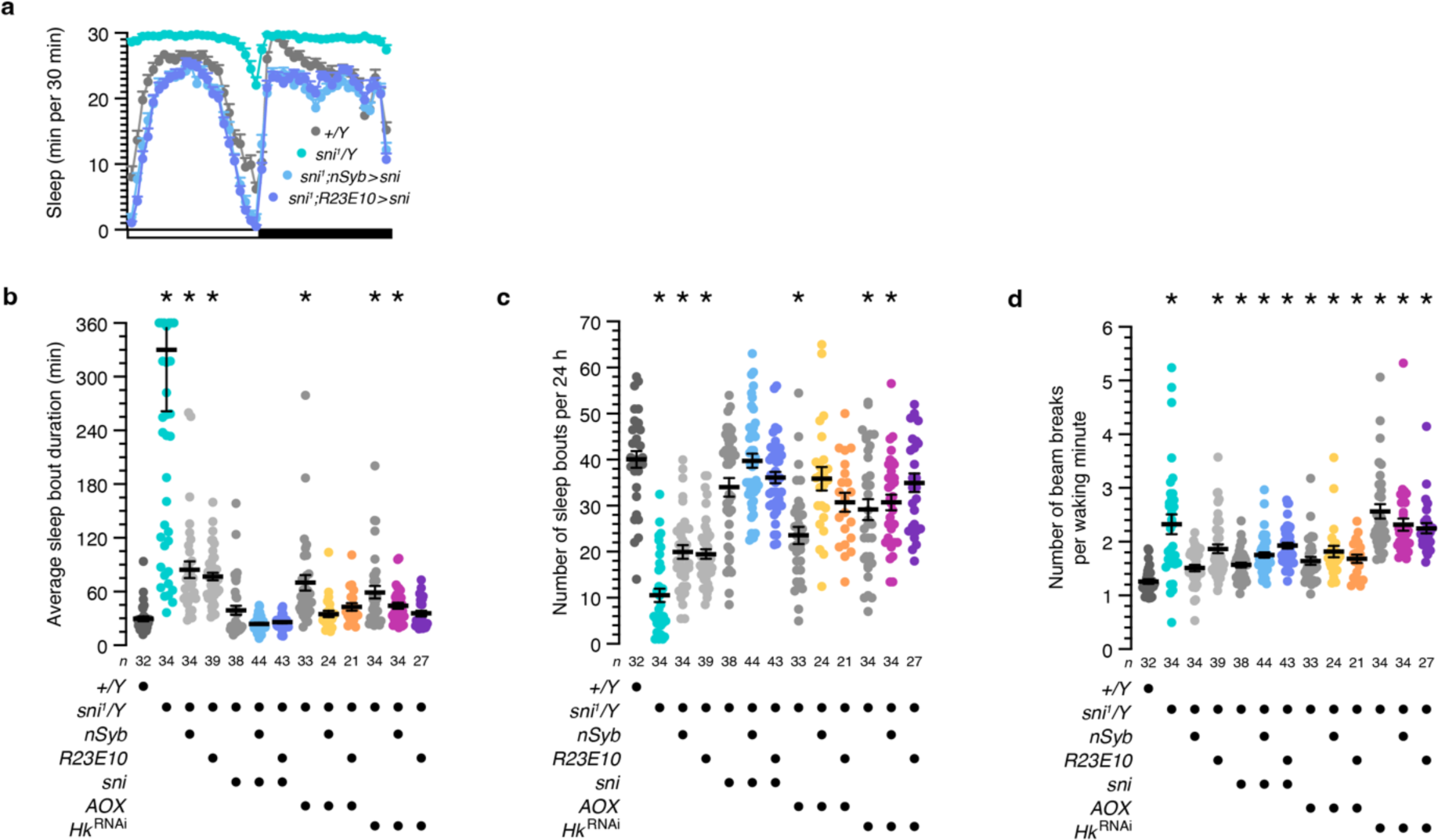
Sleep architecture and waking locomotor activity. **a**, The altered sleep profile of hemizygous males carrying the *sni^1^*allele is (over)corrected by the *nSyb-GAL4-* or *R23E10-GAL4*-driven overexpression of sniffer (*P*<0.0001 for all pairwise comparisons; two-way repeated-measures ANOVA with Holm-Šídák test; sample sizes as in **b**). **b**, The average sleep bout duration in hemizygous males carrying the *sni^1^*allele differs from wild-type (*P*<0.0001; Kruskal-Wallis ANOVA with Dunn’s test) but returns to control level if carriers also express sniffer or AOX pan-neuronally under the control of *nSyb-GAL4* (sni: *P*>0.9999; AOX: *P*>0.9999) or sniffer, AOX, or *Hk*^RNAi^ in dFBNs under the control of *R23E10-GAL4* (sni: *P*>0.9999; AOX: *P*=0.1462; *Hk*^RNAi^: *P*>0.9999). The average sleep bout durations of 6 *sni^1^* mutants exceeding 360 min are plotted at the top of the graph; mean and s.e.m. are based on the actual values. **c**, The number of sleep bouts in hemizygous males carrying the *sni^1^* allele differs from wild-type (*P*<0.0001; Kruskal-Wallis ANOVA with Dunn’s test) but returns to control level if carriers also express sniffer or AOX pan-neuronally under the control of *nSyb-GAL4* (sni: *P*>0.9999; AOX: *P*>0.9999) or sniffer, AOX, or *Hk*^RNAi^ in dFBNs under the control of *R23E10-GAL4* (sni: *P*>0.9999; AOX: *P*=0.1492; *Hk*^RNAi^: *P*>0.9999). **d**, Hemizygous males carrying the *sni^1^* allele show elevated waking locomotor activity relative to wild-type (*P*<0.0001; Kruskal-Wallis ANOVA with Dunn’s test). Asterisks indicate significant differences (*P*<0.05) from wild-type in planned pairwise comparisons. Data are means ± s.e.m. *n*, number of flies. For statistical details see Supplementary Table 2.

**Supplementary Fig. 2.**
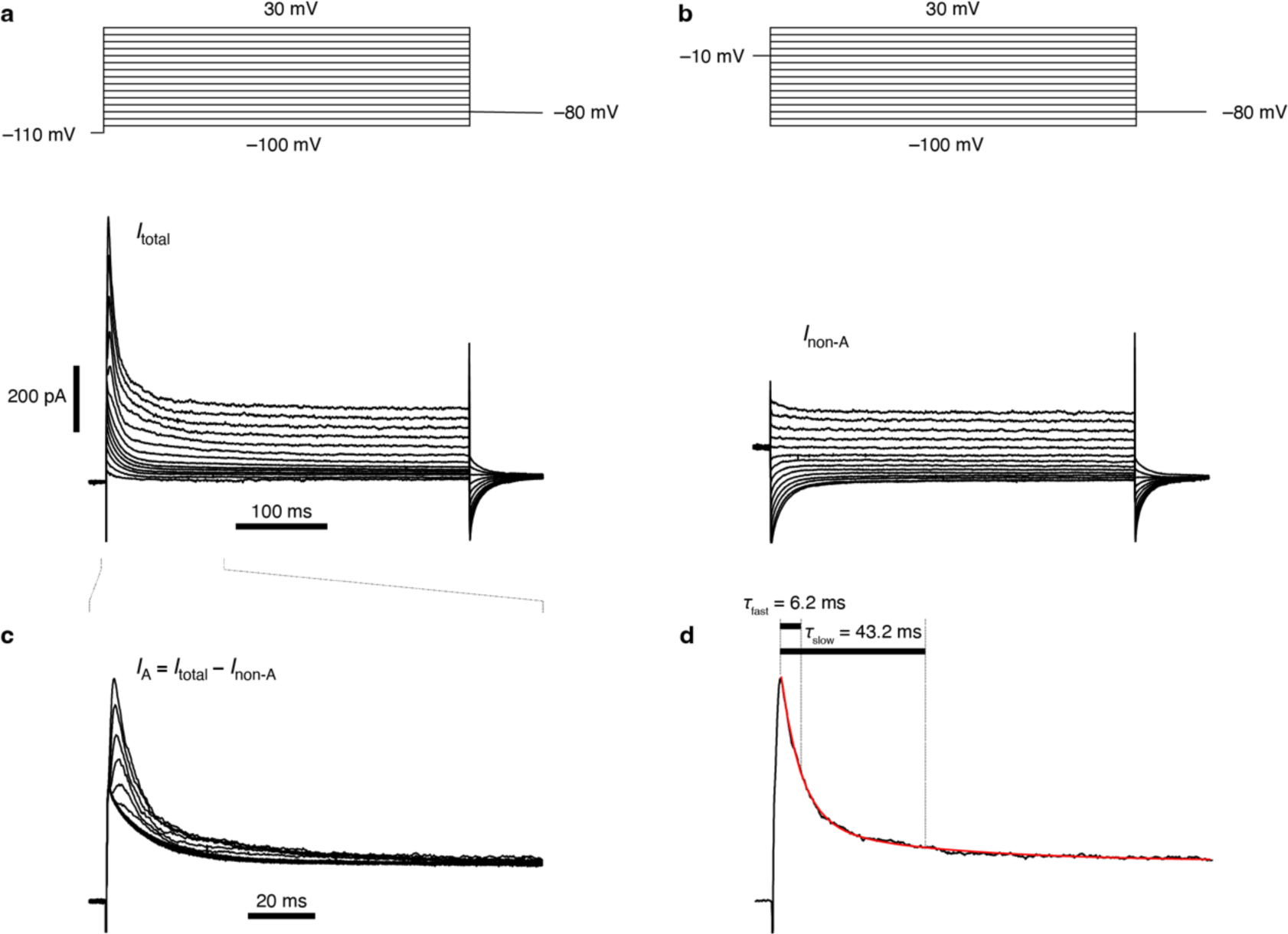
Measurement of the inactivation time constants of *I*_A_. **a**, Voltage steps from a holding potential of –110 mV (top) elicit the full complement of potassium currents in a dFBN (*I*_total_, bottom). **b**, Stepping the same neuron from a holding potential of –10 mV (top) elicits potassium currents lacking the A-type component (*I*_non-A_, bottom). **c**, Digital subtraction of *I*_non-A_ (**b**, bottom) from *I*_total_ (**a**, bottom) yields *I*_A_. Note the expanded timescale. **d**, Estimates of *τ*_fast_ and *τ*_slow_ are obtained from a double-exponential fit (red line) to the A-type current evoked by step depolarization to +30 mV.

**Supplementary Fig. 3.**
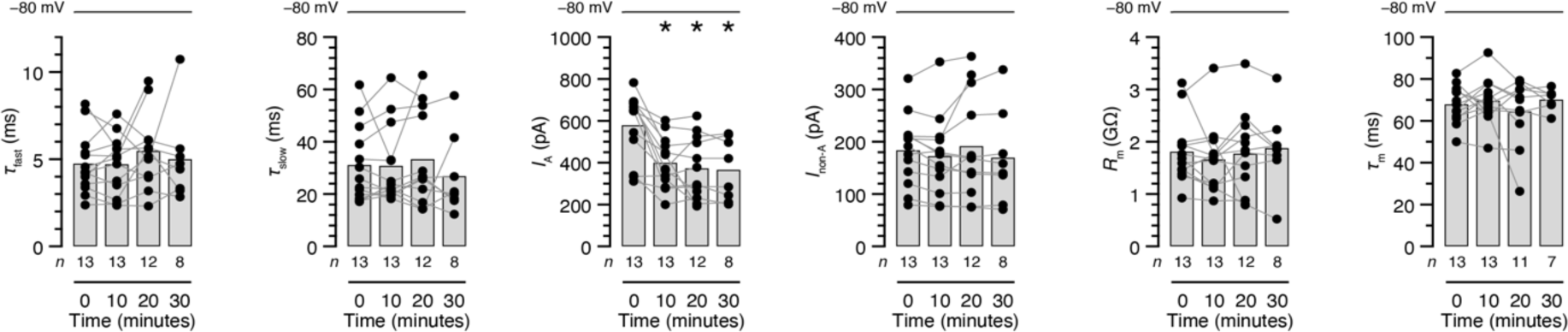
Stability of potassium currents. dFBNs were held at –80 mV, except during the voltage protocols required to measure *I*_A_. The fast and slow inactivation time constants of *I*_A_ (*τ*_fast_ and *τ*_slow_), the amplitude of *I*_non-A_, input resistances (*R*_m_), and membrane time constants (*τ*_m_) remain unchanged (*P*≥0.3527; mixed-effects model), but the amplitude of *I*_A_ runs down during the course of the recording (*P*=0.0004; mixed-effects model). Columns, population averages; dots, individual cells; *n*, number of cells; asterisks, significant differences (*P*<0.05) relative to baseline in planned comparisons by Holm-Šídák test. For statistical details see Supplementary Table 2.

**Supplementary Fig. 4.**
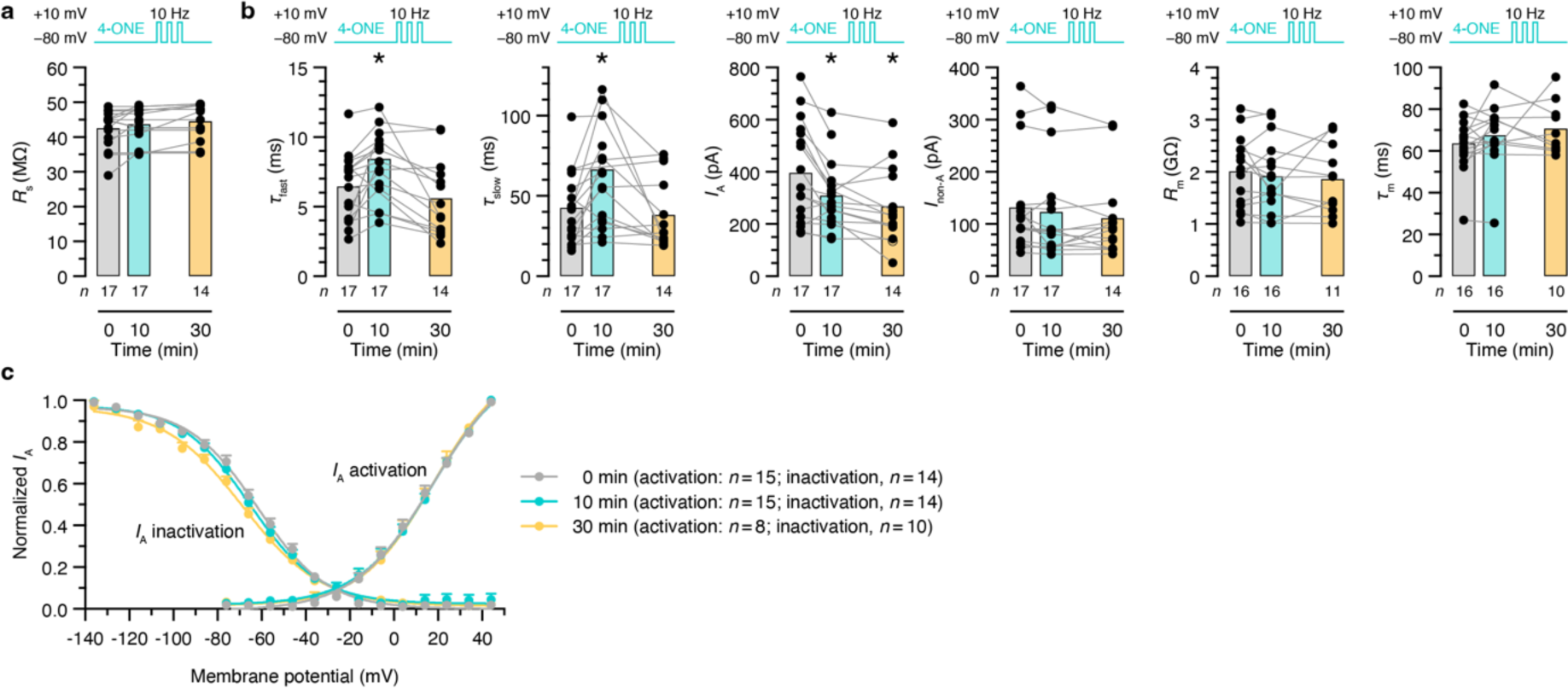
Series resistances and steady-state activation and inactivation curves. **a**, **b**, dFBNs were held at –80 mV in the interval of 0–10 minutes (except during the voltage protocols required to measure *I*_A_) and repeatedly step-depolarized to +10 mV (3 ms, 10 Hz) between 10 and 30 minutes; the intracellular solution contained 50 µM 4-ONE. Series resistances increase gradually (*R*_s_ in **a** *P*=0.0399; mixed-effects model) but remain within <20% of baseline and below 50 MΩ. The inclusion of 50 µM 4-ONE in the intracellular solution increases the fast and slow inactivation time constants of *I*_A_ above the baseline measured immediately after break-in (**b**, turquoise vs. gray shading); a series of depolarization steps between 10 and 30 minutes counteracts this increase despite the continuous presence of 4-ONE (yellow shading; *τ*_fast_: *P*=0.0054; *τ*_slow_: *P*=0.0014; mixed-effects model). The amplitude of *I*_A_ runs down during the course of the recording (*P*=0.0008; mixed-effects model); *I*_non-A_, input resistances (*R*_m_), and membrane time constants (*τ*_m_) remain unchanged (**b**, *P*≥0.2282; mixed-effects model). Columns, population averages; dots, individual cells; *n*, number of cells; asterisks, significant differences (*P*<0.05) relative to baseline in planned comparisons by Holm-Šídák test. **c**, Steady-state activation and inactivation curves of *I*_A_ in dFBNs immediately after break-in (0 minutes), after 10 minutes of dialysis with intracellular solution containing 50 µM 4-ONE (turquoise), and after a series of depolarization steps to +10 mV (3 ms, 10 Hz) between 10 and 30 minutes (yellow). Data are means ± s.e.m; solid lines, Boltzmann fits. The half-activation voltages and activation slope factors were identical at all time points (*P* = 0.5378, *F* test) but the half-inactivation voltages and inactivation slope factors differed (*P* < 0.0001, *F* test). For statistical details see Supplementary Table 2.

**Supplementary Fig. 5.**
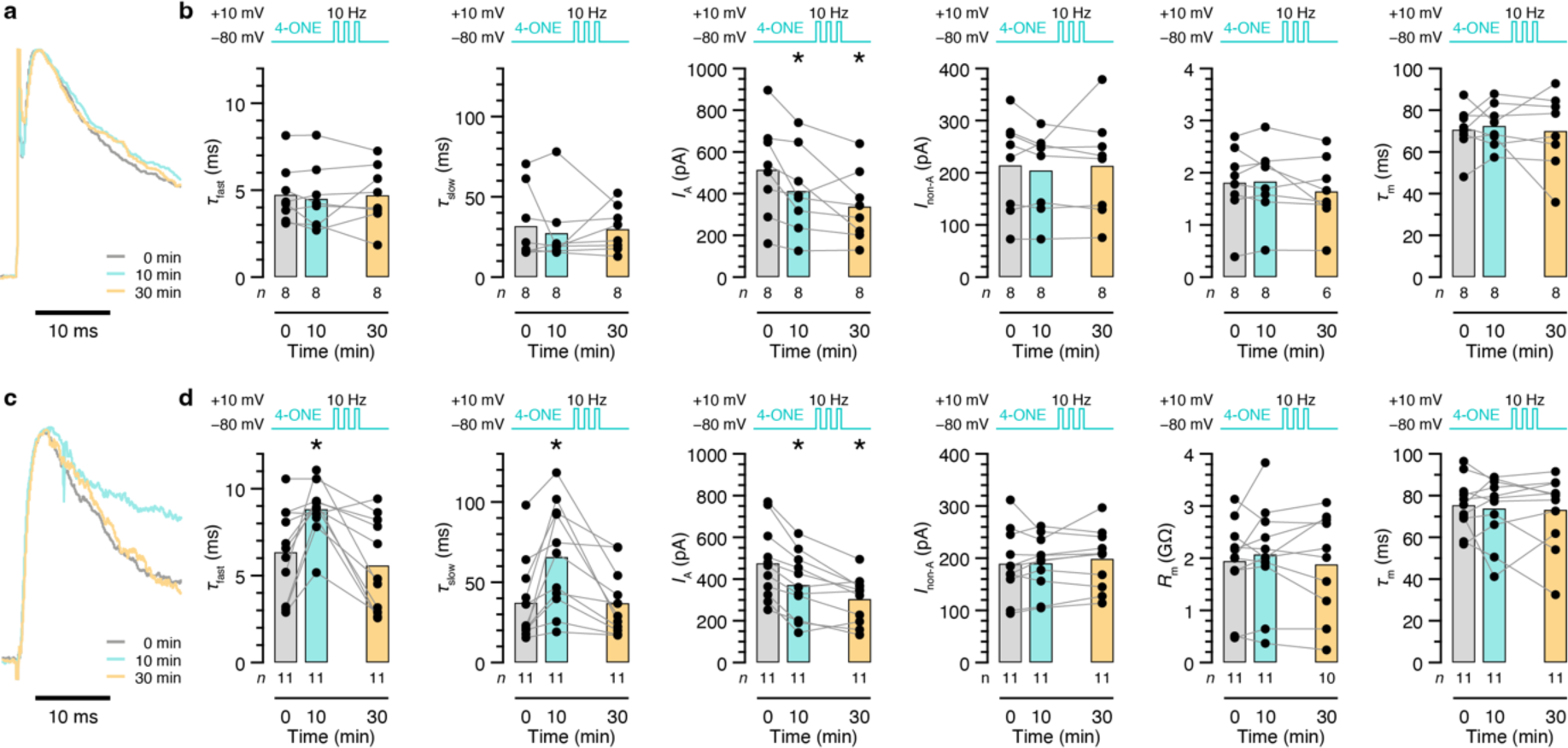
Memory storage and erasure requires a catalytically active β-subunit. **a**, **b**, dFBNs expressing a catalytically defective *Hk^K289M^*‘rescue’ transgene in a homozygous *Hk^1^* mutant background. The cells were held at –80 mV between 0 and 10 minutes (except during the voltage protocols required to measure *I*_A_) and repeatedly step-depolarized to +10 mV (3 ms, 10 Hz) between 10 and 30 minutes. The inclusion of 50 µM 4-ONE in the intracellular solution fails to increase the fast and slow inactivation time constants of *I*_A_ above the baseline measured immediately after break-in (**b**, turquoise vs. gray shading); a series of depolarization steps between 10 and 30s is similarly without effect (**b**, yellow shading; *τ*_fast_: *P*=0.6841; *τ*_slow_: *P*=0.7852; repeated-measures ANOVA; examples of peak-normalized *I*_A_ evoked in the same dFBN by voltage steps to +30 mV in **a**). The amplitude of *I*_A_ runs down during the course of the recording (**b**, *P*=0.0087; repeated-measures ANOVA); *I*_non-A_, input resistances (*R*_m_), and membrane time constants (*τ*_m_) remain unchanged (**b**, *P*≥0.2615; repeated-measures ANOVA). **c**, **d**, dFBNs expressing a catalytically competent *Hk* rescue transgene in a homozygous *Hk^1^* mutant background. The cells were held at –80 mV between 0 and 10 minutes (except during the voltage protocols required to measure *I*_A_) and repeatedly step-depolarized to +10 mV (3 ms, 10 Hz) between 10 and 30 minutes. The inclusion of 50 µM 4-ONE in the intracellular solution increases the fast and slow inactivation time constants of *I*_A_ above the baseline measured immediately after break-in (**d**, turquoise vs. gray shading); a series of depolarization steps between 10 and 30 s counteracts this increase despite the continuous presence of 4-ONE (**d**, yellow shading; *τ*_fast_: *P*=0.0005; *τ*_slow_: *P*=0.0012; repeated-measures ANOVA; examples of peak-normalized *I*_A_ evoked in the same dFBN by voltage steps to +30 mV in **c**). The amplitude of *I*_A_ runs down during the course of the recording (**d**, *P*<0.0001; repeated-measures ANOVA); *I*_non-A_, input resistances (*R*_m_), and membrane time constants (*τ*_m_) remain unchanged (**d**, *P*≥0.4334; repeated-measures ANOVA or mixed-effects model). Columns, population averages; dots, individual cells; *n*, number of cells; asterisks, significant differences (*P*<0.05) relative to baseline in planned comparisons by Holm-Šídák test. For statistical details see Supplementary Table 2.

**Supplementary Fig. 6.**
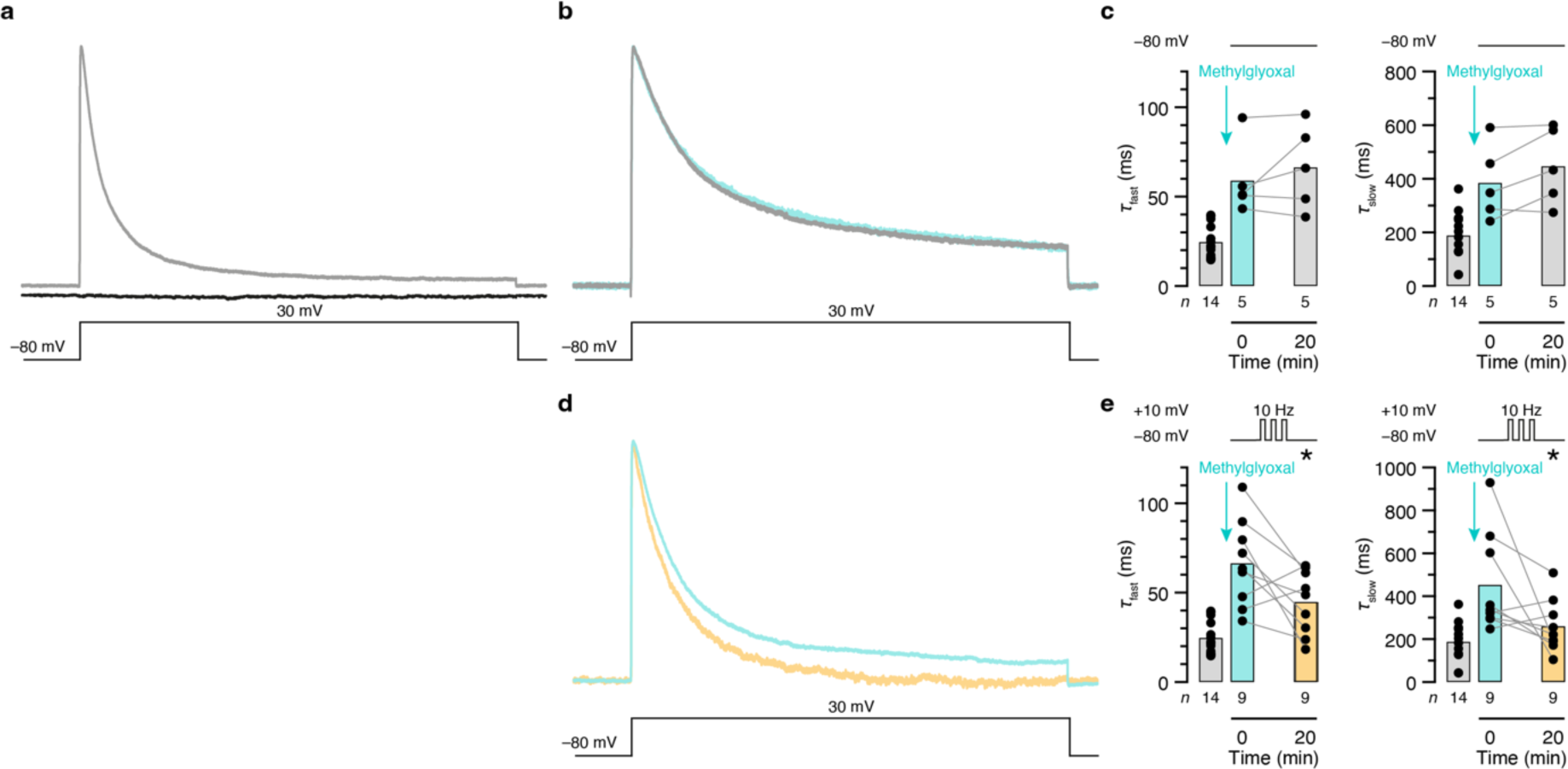
Memory storage and erasure by mammalian K_V_1.4–β2 channels in a heterologous expression system. **a**, Examples of peak-normalized transmembrane currents evoked by 1-s voltage pulses from –80 mV to +30 mV in HEK-293 cells expressing mouse K_V_1.4 and K_V_β2 (gray trace), or in untransfected HEK-293 cells (black trace). **b**–**e**, HEK-293 cells expressing mouse K_V_1.4 and K_V_β2. A 1-h exposure to 12 mM methylglyoxal, followed by three washes with methylglyoxal-free solution, increases the fast and slow inactivation time constants of transmembrane currents relative to those of cells maintained in the absence of methylglyoxal (**c**, **e**, turquoise vs. gray shading; *τ*_fast_: *P*<0.0001; *τ*_slow_: *P*<0.0001; Mann-Whitney test). In cells held at –80 mV (except during the voltage protocols required to measure *I*_A_), the time constants remain stably elevated for 20 minutes **(c**, *τ*_fast_: *P*=0.4375; *τ*_slow_: *P*=0.1875; Wilcoxon test; examples of peak-normalized currents in **b**), but a series of depolarization steps (3 ms, 10 Hz, 20 minutes) to +10 mV reverses the increase (**e**, yellow shading; *τ*_fast_: *P*=0.0370, paired *t*-test; *τ*_slow_: *P*=0.0391, Wilcoxon test; examples of peak-normalized currents in **d**). Columns, population averages; dots, individual cells; *n*, number of cells; asterisks, significant differences (*P*<0.05) relative to the 0-minute time point. For statistical details see Supplementary Table 2.

**Supplementary Fig. 7.**
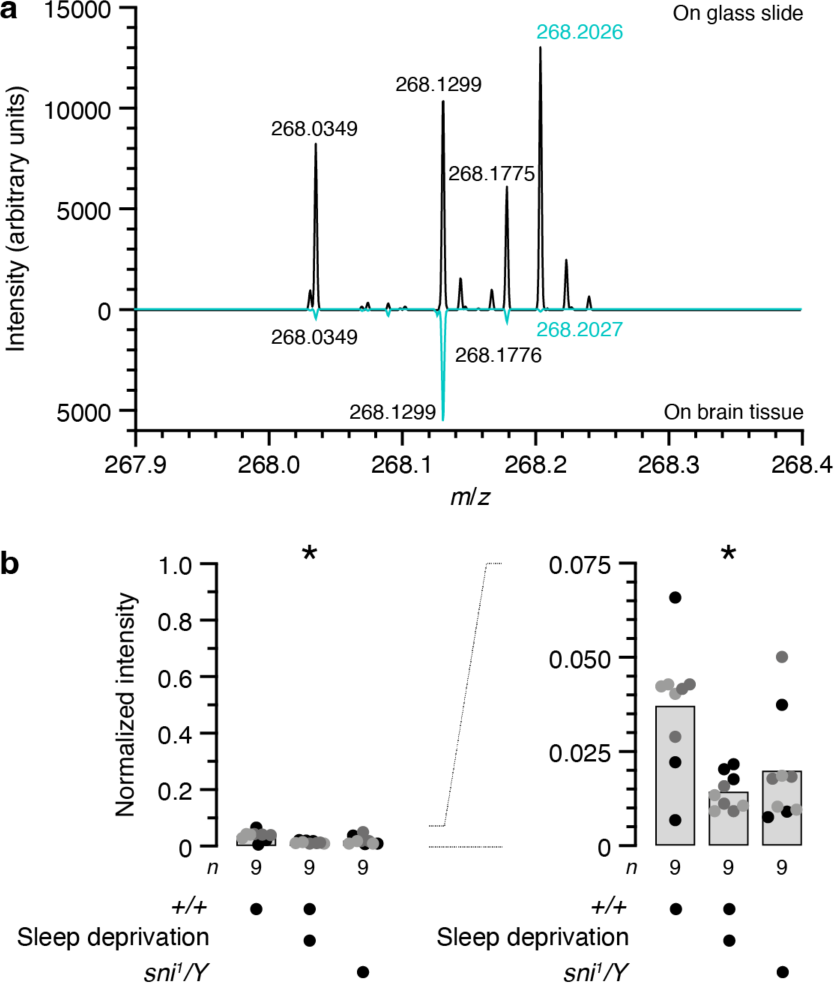
SMALDI-MSI analysis of 4-ONE. **a**, Mirror plot of mass spectra of 100 µM 4-ONE standard on a blank slide (top) or a brain cryosection (bottom). Spectra were acquired in single-ion-monitoring mode at the calculated *m*/*z* of the [4-ONE+GirT-H_2_O]^+^ ion (268.2020); peaks with a mass deviation <5 ppm are labelled in green type. **b**, The intensity of the [4-ONE+GirT-H_2_O]^+^ signal is decreased in cryosections of sleep-deprived brains (*P*=0.0179, Kruskal-Wallis ANOVA) but not significantly altered in hemizygous males carrying the *sni^1^* allele (*P*=0.0560). Intensities on the left are normalized to a 100 µM 4-ONE standard on a blank slide; the scale is expanded on the right. Columns, population averages; dots, technical replicates of 3 biological replicates grouped by shades of gray; *n*, number of replicates; asterisks, significant differences (*P*<0.05) relative to rested wild-type flies in planned comparisons by Dunnett’s T3 test. For statistical details see Supplementary Table 2.

**Supplementary Table 1.**
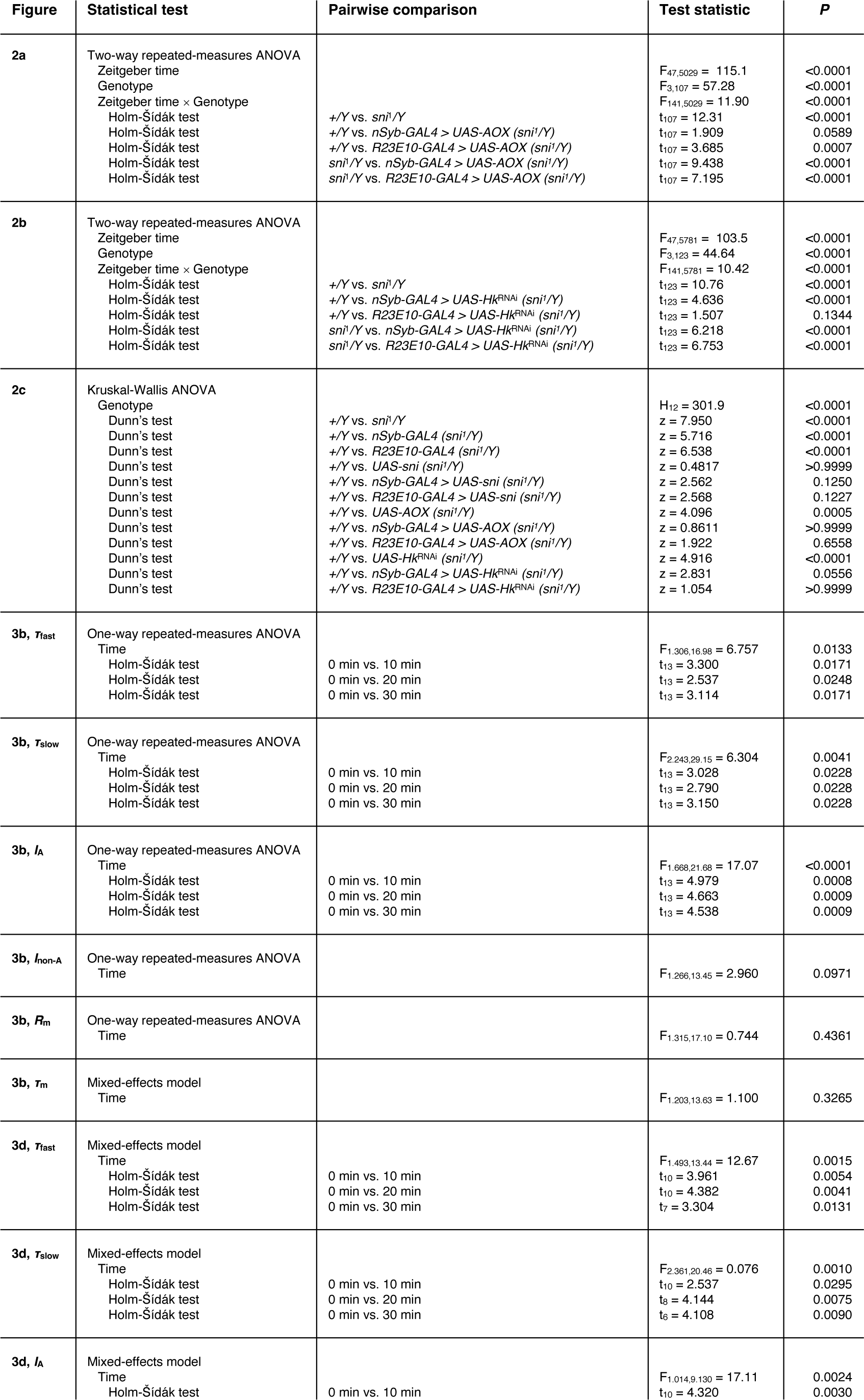
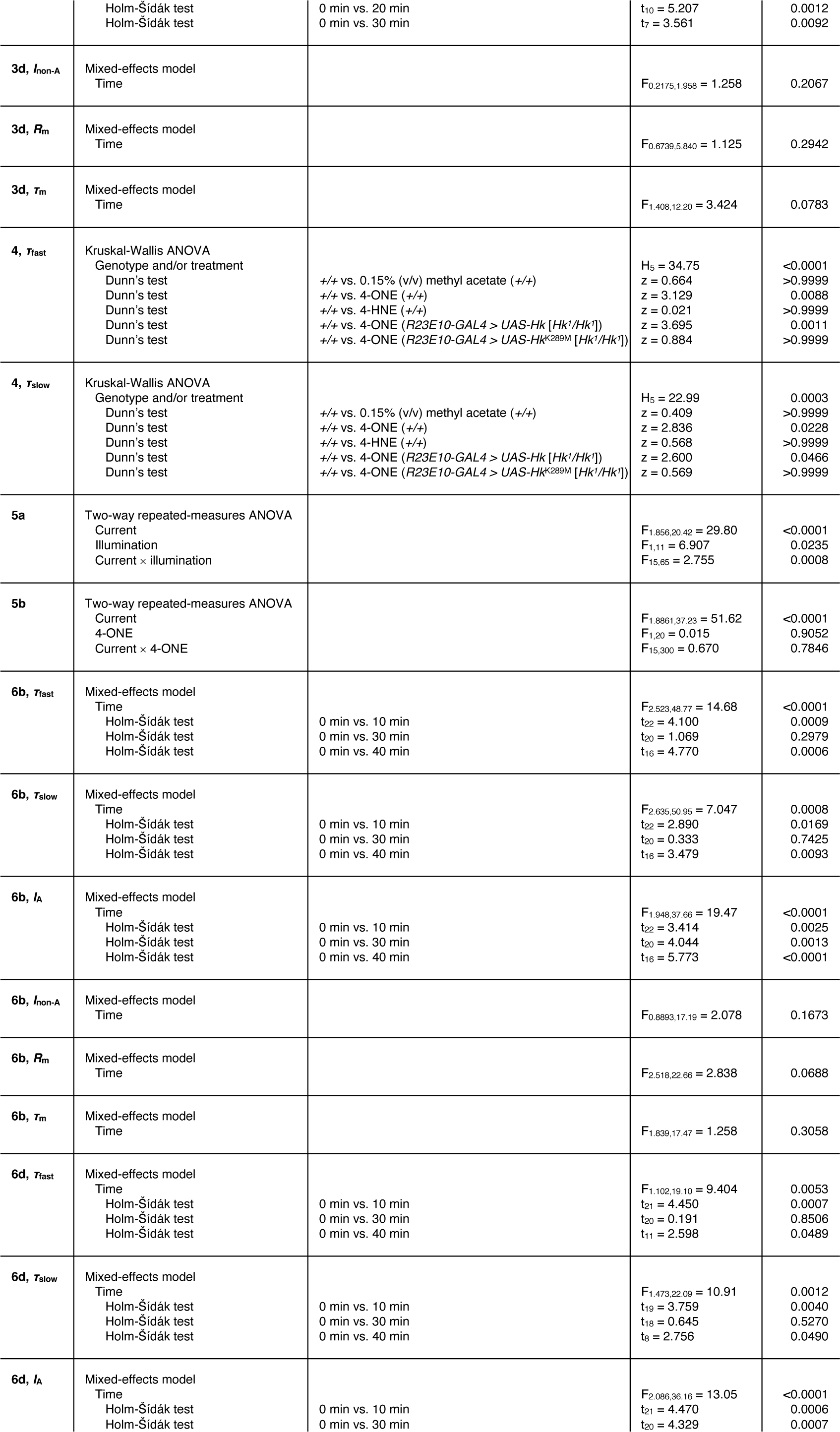
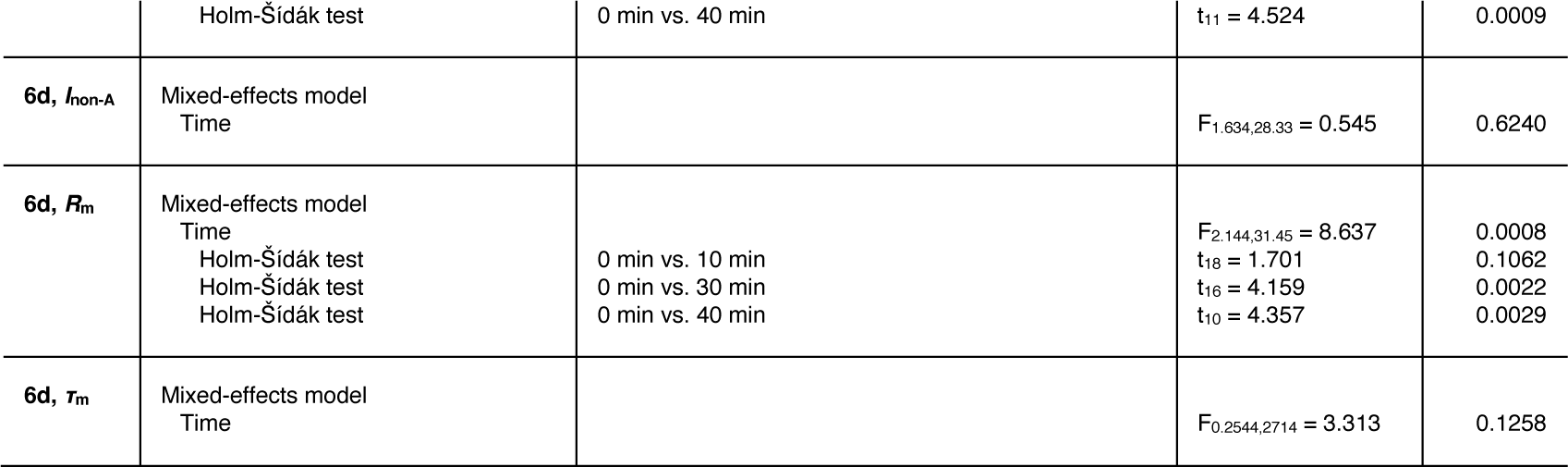
Statistical analyses of Figures 2–6.

**Supplementary Table 2.**
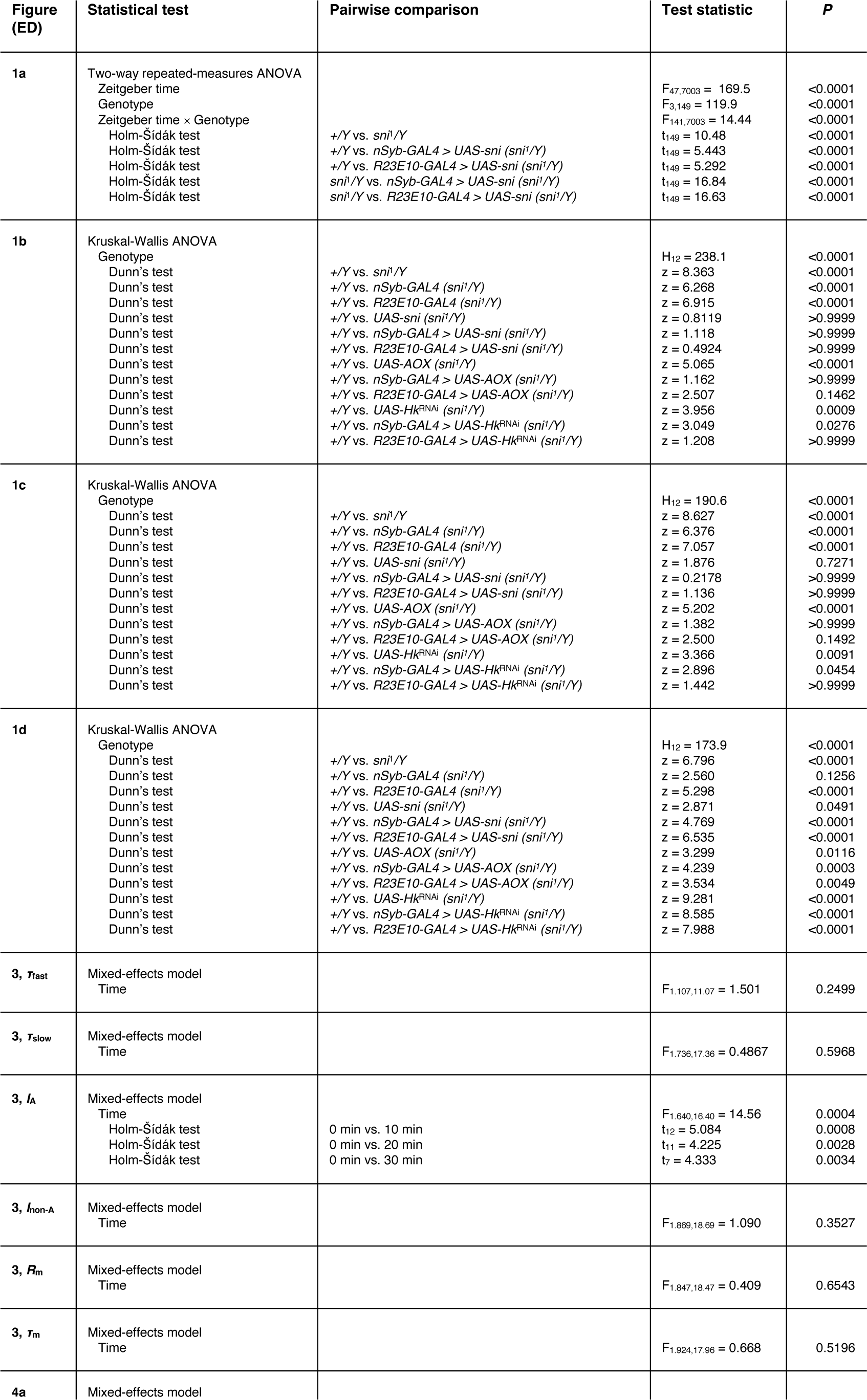
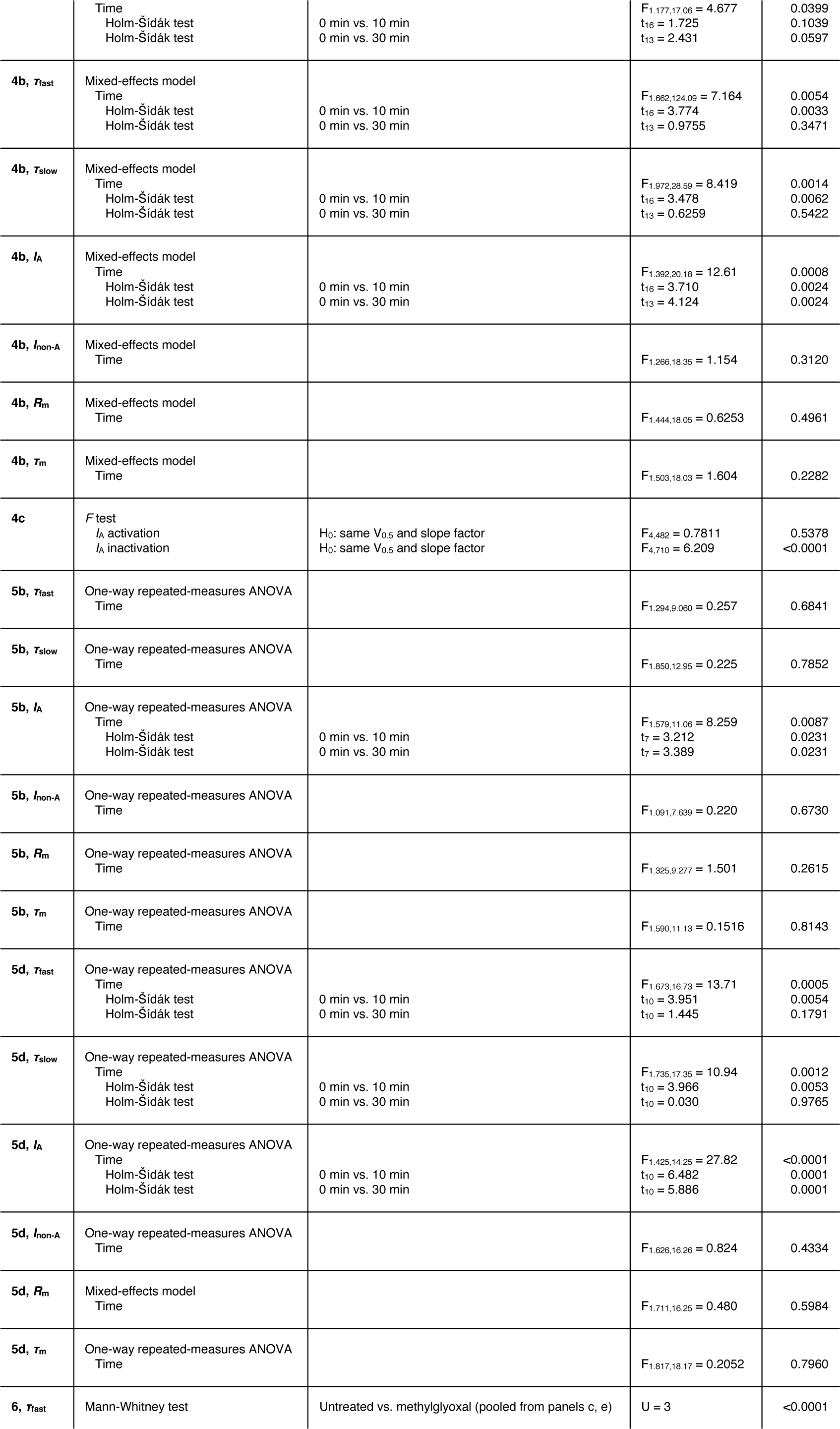
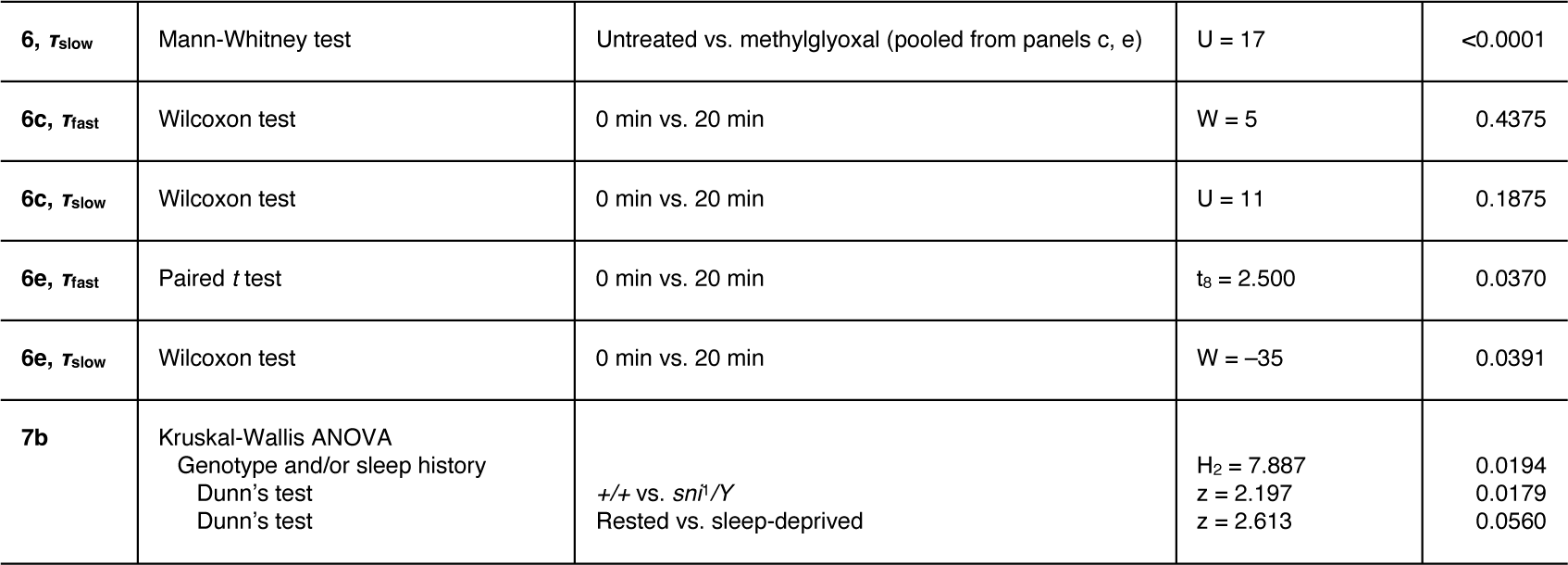
Statistical analyses of Extended Data Figures 1–7.

## References

1. Rehm, H. & Lazdunski, M. Purification and subunit structure of a putative K+-channel protein identified by its binding properties for dendrotoxin I. Proc Natl Acad Sci U S A 85, 4919–4923 (1988).

2. Parcej, D. N., Scott, V. E. & Dolly, J. O. Oligomeric properties of α-dendrotoxin-sensitive potassium ion channels purified from bovine brain. Biochemistry 31, 11084–11088 (1992).

3. Nakahira, K., Shi, G., Rhodes, K. J. & Trimmer, J. S. Selective interaction of voltage-gated K+ channel β-subunits with α-subunits. J Biol Chem 271, 7084–7089 (1996).

4. Long, S. B., Campbell, E. B. & MacKinnon, R. Crystal structure of a mammalian voltage-dependent Shaker family K+ channel. Science 309, 897–903 (2005).

5. Pongs, O. & Schwarz, J. R. Ancillary subunits associated with voltage-dependent K+ channels. Physiol Rev 90, 755–796 (2010).

6. McCormack, T. & McCormack, K. Shaker K+ channel β subunits belong to an NAD(P)H-dependent oxidoreductase superfamily. Cell 79, 1133–1135 (1994).

7. Chouinard, S. W., Wilson, G. F., Schlimgen, A. K. & Ganetzky, B. A potassium channel β subunit related to the aldo-keto reductase superfamily is encoded by the Drosophila Hyperkinetic locus. Proc Natl Acad Sci U S A 92, 6763–6767 (1995).

8. Weng, J., Cao, Y., Moss, N. & Zhou, M. Modulation of voltage-dependent Shaker family potassium channels by an aldo-keto reductase. J Biol Chem 281, 15194–15200 (2006).

9. Tipparaju, S. M., Barski, O. A., Srivastava, S. & Bhatnagar, A. Catalytic mechanism and substrate specificity of the β-subunit of the voltage-gated potassium channel. Biochemistry 47, 8840–8854 (2008).

10. Gulbis, J. M., Mann, S. & MacKinnon, R. Structure of a voltage-dependent K+ channel β subunit. Cell 97, 943–952 (1999).

11. Tempel, B. L., Papazian, D. M., Schwarz, T. L., Jan, Y. N. & Jan, L. Y. Sequence of a probable potassium channel component encoded at Shaker locus of Drosophila. Science 237, 770–775 (1987).

12. Iverson, L. E., Tanouye, M. A., Lester, H. A., Davidson, N. & Rudy, B. A-type potassium channels expressed from Shaker locus cDNA. Proc Natl Acad Sci U S A 85, 5723–5727 (1988).

13. Cirelli, C. et al. Reduced sleep in Drosophila Shaker mutants. Nature 434, 1087–1092 (2005).

14. Bushey, D., Huber, R., Tononi, G. & Cirelli, C. Drosophila Hyperkinetic mutants have reduced sleep and impaired memory. J Neurosci 27, 5384–5393 (2007).

15. Donlea, J. M., Thimgan, M. S., Suzuki, Y., Gottschalk, L. & Shaw, P. J. Inducing sleep by remote control facilitates memory consolidation in Drosophila. Science 332, 1571–1576 (2011).

16. Pimentel, D. et al. Operation of a homeostatic sleep switch. Nature 536, 333–337 (2016).

17. Donlea, J. M., Pimentel, D. & Miesenböck, G. Neuronal machinery of sleep homeostasis in Drosophila. Neuron 81, 860–872 (2014).

18. Kempf, A., Song, S. M., Talbot, C. B. & Miesenböck, G. A potassium channel β-subunit couples mitochondrial electron transport to sleep. Nature 568, 230–234 (2019).

19. Bähring, R. et al. Coupling of voltage-dependent potassium channel inactivation and oxidoreductase active site of Kvβ subunits. J Biol Chem 276, 22923–22929 (2001).

20. Pan, Y., Weng, J., Cao, Y., Bhosle, R. C. & Zhou, M. Functional coupling between the Kv1.1 channel and aldoketoreductase Kvβ1. J Biol Chem 283, 8634–8642 (2008).

21. Chintaluri, C. & Vogels, T. P. Metabolically regulated spiking could serve neuronal energy homeostasis and protect from reactive oxygen species. Proc Natl Acad Sci U S A 120, e2306525120 (2023).

22. Hasenhuetl, P. S. et al. A half-centre oscillator encodes sleep pressure. Submitted. (2024).

23. Yin, H., Xu, L. & Porter, N. A. Free radical lipid peroxidation: Mechanisms and analysis. Chem Rev 111, 5944–5972 (2011).

24. Ayala, A., Muñoz, M. F. & Argüelles, S. Lipid peroxidation: Production, metabolism, and signaling mechanisms of malondialdehyde and 4-hydroxy-2-nonenal. Oxid Med Cell Longev 2014, 360438 (2014).

25. Willems, P. H., Rossignol, R., Dieteren, C. E., Murphy, M. P. & Koopman, W. J. Redox homeostasis and mitochondrial dynamics. Cell Metab 22, 207–218 (2015).

26. Esterbauer, H., Schaur, R. J. & Zollner, H. Chemistry and biochemistry of 4-hydroxynonenal, malonaldehyde and related aldehydes. Free Radic Biol Med 11, 81–128 (1991).

27. Parvez, S., Long, M. J. C., Poganik, J. R. & Aye, Y. Redox signaling by reactive electrophiles and oxidants. Chem Rev 118, 8798–8888 (2018).

28. Dennard, R. H. How we made DRAM. Nat Electron 1, 372 (2018).

29. Vaccaro, A. et al. Sleep loss can cause death through accumulation of reactive oxygen species in the gut. Cell 181, 1307–1328.e15 (2020).

30. Kent, C. Eukaryotic phospholipid biosynthesis. Annu Rev Biochem 64, 315–343 (1995).

31. Vance, J. E. Phospholipid synthesis and transport in mammalian cells. Traffic 16, 1–18 (2015).

32. Choi, S. Y. et al. A common lipid links Mfn-mediated mitochondrial fusion and SNARE-regulated exocytosis. Nat Cell Biol 8, 1255–1262 (2006).

33. Sarnataro, R., Velasco, C. D., Monaco, N., Kempf, A. & Miesenböck, G. Mitochondrial origins of the pressure to sleep. Submitted. (2024).

34. Wai, T. & Langer, T. Mitochondrial dynamics and metabolic regulation. Trends Endocrinol Metab 27, 105–117 (2016).

35. Fridovich, I. Oxygen toxicity: A radical explanation. J Exp Biol 201, 1203–1209 (1998).

36. Doorn, J. A., Maser, E., Blum, A., Claffey, D. J. & Petersen, D. R. Human carbonyl reductase catalyzes reduction of 4-oxonon-2-enal. Biochemistry 43, 13106–13114 (2004).

37. Oppermann, U. Carbonyl reductases: The complex relationships of mammalian carbonyl-and quinone-reducing enzymes and their role in physiology. Annu Rev Pharmacol Toxicol 47, 293–322 (2007).

38. Botella, J. A. et al. The Drosophila carbonyl reductase sniffer prevents oxidative stress-induced neurodegeneration. Curr Biol 14, 782–786 (2004).

39. Martin, H. J. et al. The Drosophila carbonyl reductase sniffer is an efficient 4-oxonon-2-enal (4ONE) reductase. Chem Biol Interact 191, 48–54 (2011).

40. Hill, V. M. et al. A bidirectional relationship between sleep and oxidative stress in Drosophila. PLoS Biol 16, e2005206 (2018).

41. Maxwell, D. P., Wang, Y. & McIntosh, L. The alternative oxidase lowers mitochondrial reactive oxygen production in plant cells. Proc Natl Acad Sci U S A 96, 8271–8276 (1999).

42. Shu, X. et al. A genetically encoded tag for correlated light and electron microscopy of intact cells, tissues, and organisms. PLoS Biol 9, e1001041 (2011).

43. Murphy, M. P. et al. Guidelines for measuring reactive oxygen species and oxidative damage in cells and in vivo. Nat Metab 4, 651–662 (2022).

44. Hattori, S., Murakami, F. & Song, W. J. Rundown of a transient potassium current is attributable to changes in channel voltage dependence. Synapse 48, 57–65 (2003).

45. Fogle, K. J. et al. CRYPTOCHROME-mediated phototransduction by modulation of the potassium ion channel β-subunit redox sensor. Proc Natl Acad Sci U S A 112, 2245–2250 (2015).

46. Bar-Yehuda, D. & Korngreen, A. Space-clamp problems when voltage clamping neurons expressing voltage-gated conductances. J Neurophysiol 99, 1127–1136 (2008).

47. Sallin, O. et al. Semisynthetic biosensors for mapping cellular concentrations of nicotinamide adenine dinucleotides. Elife 7, e32638 (2018).

48. Gopalakrishnan, A., Ji, L. L. & Cirelli, C. Sleep deprivation and cellular responses to oxidative stress. Sleep 27, 27–35 (2004).

49. Silva, R. H. et al. Role of hippocampal oxidative stress in memory deficits induced by sleep deprivation in mice. Neuropharmacology 46, 895–903 (2004).

50. Bellesi, M. et al. Sleep loss promotes astrocytic phagocytosis and microglial activation in mouse cerebral cortex. J Neurosci 37, 5263–5273 (2017).

51. Kaplan, W. D. & Trout, W. E. The behavior of four neurological mutants of Drosophila. Genetics 61, 399–409 (1969).

52. Stern, M. & Ganetzky, B. Altered synaptic transmission in Drosophila Hyperkinetic mutants. J Neurogenet 5, 215–228 (1989).

53. Jenett, A. et al. A GAL4-driver line resource for Drosophila neurobiology. Cell Rep 2, 991–1001 (2012).

54. Ng, J. et al. Genetically targeted 3D visualisation of Drosophila neurons under electron microscopy and x-ray microscopy using miniSOG. Sci Rep 6, 38863 (2016).

55. Fernandez-Ayala, D. J. M. et al. Expression of the Ciona intestinalis alternative oxidase (AOX) in Drosophila complements defects in mitochondrial oxidative phosphorylation. Cell Metab 9, 449–460 (2009).

56. Dietzl, G. et al. A genome-wide transgenic RNAi library for conditional gene inactivation in Drosophila. Nature 448, 151–156 (2007).

57. Simpson, J. H. Rationally subdividing the fly nervous system with versatile expression reagents. J Neurogenet 30, 185–194 (2016).

58. Hendricks, J. C. et al. Rest in Drosophila is a sleep-like state. Neuron 25, 129–138 (2000).

59. Shaw, P. J., Cirelli, C., Greenspan, R. J. & Tononi, G. Correlates of sleep and waking in Drosophila melanogaster. Science 287, 1834–1837 (2000).

60. Vecsey, C. G., Koochagian, C., Porter, M. T., Roman, G. & Sitaraman, D. Analysis of sleep and circadian rhythms from Drosophila Activity-Monitoring data using SCAMP. Cold Spring Harb Protoc (2024).

61. Shaw, P. J., Tononi, G., Greenspan, R. J. & Robinson, D. F. Stress response genes protect against lethal effects of sleep deprivation in Drosophila. Nature 417, 287–291 (2002).

62. Spengler, B. & Hubert, M. Scanning microprobe matrix-assisted laser desorption ionization (SMALDI) mass spectrometry: Instrumentation for sub-micrometer resolved LDI and MALDI surface analysis. J Am Soc Mass Spectrom 13, 735–748 (2002).

63. Dreisbach, D., Heiles, S., Bhandari, D. R., Petschenka, G. & Spengler, B. Molecular networking and on-tissue chemical derivatization for enhanced identification and visualization of steroid glycosides by MALDI mass spectrometry imaging. Anal Chem 94, 15971–15979 (2022).

64. Paschke, C. et al. Mirion—a software package for automatic processing of mass spectrometric images. J Am Soc Mass Spectrom 24, 1296–1306 (2013).

65. Müller, M. A., Kompauer, M., Strupat, K., Heiles, S. & Spengler, B. Implementation of a high-repetition-rate laser in an AP-SMALDI MSI system for enhanced measurement performance. J Am Soc Mass Spectrom 32, 465–472 (2021).

66. Sud, M. et al. LMSD: LIPID MAPS structure database. Nucleic Acids Res 35, D527–32 (2007).

67. Koelmel, J. P. et al. LipidMatch: An automated workflow for rule-based lipid identification using untargeted high-resolution tandem mass spectrometry data. BMC Bioinformatics 18, 331 (2017).

68. Neher, E. Correction for liquid junction potentials in patch clamp experiments. Methods Enzymol 207, 123–131 (1992).

69. Ruppersberg, J. P. et al. Regulation of fast inactivation of cloned mammalian IK(a) channels by cysteine oxidation. Nature 352, 711–714 (1991).

70. Lin, W. J. et al. LipidSig: A web-based tool for lipidomic data analysis. Nucleic Acids Res 49, W336–W345 (2021).

